# Developmental regulation of progenitor aging shapes long-term intestinal homeostasis in *Drosophila*

**DOI:** 10.64898/2026.03.21.713357

**Authors:** Shikha Malik, Atharva Anand Mahajan, Saraswathi J Pillai, Isha Shinde, Muhamad Shameem, Pratik Chandrani, Mandar M Inamdar, Rohan J Khadilkar

## Abstract

Aging causes a progressive loss of tissue homeostasis, with stem cell exhaustion as a major hallmark. Age-associated decline in organ function is widely perceived as emanating from progressive accumulation of cellular damage in adult tissues. However, whether aging trajectories are established early on during development remains an open question. Here, we demonstrate that genetic modulation of cellular aging pathways in larval adult midgut progenitors (AMPs), which serve as the precursors of adult intestinal stem cells and differentiated epithelial cells, dictates the long-term trajectory of intestinal aging in *Drosophila*. Accelerated cellular aging by genetic perturbation employing Toll or Imd pathway overactivation or elevation of reactive oxygen species (ROS) using ND42 (mitochondrial complex I) knockdown in the AMPs results in aberrant progenitor proliferation, skewed lineage allocation, epithelial barrier dysfunction, and genomic instability. These alterations are accompanied by marked destabilization of AMP islet architecture and widespread changes in age-related molecular signatures, as revealed by bulk transcriptomic analysis. In contrast, decelerated cellular aging mediated by Foxo or Atg8a overexpression results in a decrease in enteroendocrine population and the intestinal barrier remained unaffected. Intriguingly, early-life activation of immune and oxidative stress signaling manifested later in the adult gut as elevated enteroendocrine differentiation, highlighting lasting effects on intestinal regenerative capacity and lineage balance. Together, our findings demonstrate that cellular aging is tightly regulated early on in development and its perturbation can cause developmental disruption hampering adult gut homeostasis, establishing AMPs as key developmental determinants that regulate the trajectory of intestinal aging in *Drosophila*.

## Introduction

Aging is a dynamic and multifaceted biological process marked by a gradual deterioration of cellular and tissue homeostasis, ultimately compromising organismal function [1, 2]. Such age-driven decline is the principal risk factor underlying most chronic and degenerative diseases like cancer, diabetes, cardiovascular, neurodegenerative [2, 3], and metabolic disorders [4]. Recent aging paradigms have highlighted the core hallmarks of aging as genomic instability, loss of proteostasis, deregulated nutrient sensing, cellular senescence, stem cell exhaustion, etc. [2, 5]. Notably, these hallmarks are functionally interconnected, whereby perturbation of one often triggers alterations in another [2].

Among these hallmarks, stem cell exhaustion has emerged as a pivotal driver of age-associated tissue deterioration, particularly in tissues with high turnover rate [6]. Aging impairs the ability of stem cell regeneration, self-renewal, tissue development, and lineage commitment across various tissues [7]. For instance, aged hematopoietic stem cells exhibit increased clonal hematopoiesis, impaired regenerative potential and elevated myelopoiesis [8–10]; neural stem cells display diminished potential for self-renewal and differentiation, altered protein homeostasis, and mitochondrial dysfunction with age [7, 11, 12]; muscle stem cells aging is accompanied by decreased regenerative capacity, reduced satellite cell pool, impaired autophagic activity and mitochondrial dysfunction [13–15]. Likewise, mammalian intestinal stem cells undergo age-related functional decline marked by skewed lineage differentiation, reduced regenerative potential, compromised barrier integrity, increased inflammation [7, 16, 17] and reduced canonical Wnt and Notch signaling activity [18].

*Drosophila* serves as a genetically tractable model to study conserved longevity and stress-associated pathways [4, 19–21], with the *Drosophila* intestine serving as a well-established stem-cell driven system to investigate aging and organ homeostasis [22–24]. During *Drosophila* embryogenesis, adult midgut progenitors (AMPs) arise in the midgut from an intermediate endoblast population and persist as undifferentiated, multipotent progenitors throughout the larval development [25]. During early larval stages, AMPs proliferate extensively and disperse along the midgut epithelium before forming discrete clusters, known as midgut imaginal islets, by the third instar stage [26, 27]. These proliferating AMP clusters are encased by one or more peripheral cells that function as a temporary niche, which delays differentiation until the onset of metamorphosis [28]. During metamorphosis, the larval gut undergoes histolysis via programmed cell death, and a new adult midgut epithelium is generated from these AMP clusters [27, 29], giving rise to adult intestinal stem cells (ISCs), absorptive enterocytes (ECs), and secretory enteroendocrine (EEs) cells.

In adult flies, aging is accompanied by epithelial dysfunction, manifested as altered cell populations with increased ISCs, and EEs, reduced EC numbers, compromised barrier, and tissue integrity [1, 30–35], and microbial dysbiosis [36, 37]. Despite these insights, it remains unclear whether age-related perturbations in progenitor population early on in development can shape organ homeostasis in adults for example whether perturbations in adult midgut progenitors (AMPs) in the larvae shape intestinal homeostasis during development and in the adult gut. Given the precise regulation of AMP proliferation and lineage specification during development, these progenitors may represent the critical window of vulnerability, wherein early-life perturbation or stress predisposes the intestine to age-associated functional decline.

Aging in *Drosophila* is driven by the synergistic interplay between inflammaging and oxidative stress, two interrelated processes that gradually destabilize cellular and tissue homeostasis [38–40]. Age-associated chronic inflammation, termed inflammaging, has been recognized as a new hallmark of aging and is marked by chronic low-grade immune activation that promotes tissue degeneration [2, 41, 42]. In *Drosophila*, inflammaging stems from immunosenescence, wherein consistent activation of NF-κB pathways like IMD and Toll signaling converts acute protective responses into chronic, tissue-damaging inflammation [43–47]. Sustained immune activation disrupts stem cell function by elevating antimicrobial peptide and pro-inflammatory cytokine expression in *Drosophila*, thereby reducing lifespan, and promoting tissue regeneration [3, 40, 48]. Parallelly, aging also triggers an imbalance between mitochondrial reactive oxygen species (ROS) production and antioxidant defenses like superoxide dismutase 1 (SOD1), causing oxidative damage to biomolecules, further reinforcing the inflammatory cascade [49–53]. Accordingly, disruption of ROS homeostasis has been shown to accelerate sperm aging in *Drosophila*, underscoring the importance of redox balance in longevity and tissue integrity [54]. This stochastic nature of events forms a feedback loop where inflammatory signaling and oxidative damage mutually accelerate cellular senescence and organismal aging. On the contrary, Foxo transcription factors (such as *dFOXO* in flies or *FoxO3* in humans) serve as master regulators of longevity by upregulating genes for stress resistance, DNA repair, and ROS scavengers like super oxide dismutase 2 (SOD2) and catalase [55, 56]. In *Drosophila*, Foxo activation enhances oxidative stress resistance and proteostasis to promote organismal lifespan, highlighting Foxo’s conserved role in tissue homeostasis [57, 58]. Complementing this, autophagy acts as a vital housekeeping process that maintains cellular homeostasis by utilizing lysosomal degradation to recycle damaged organelles and proteins; however, this flux naturally declines with age [59–62]. Enhancing autophagy, specifically through overactivation of Atg8a or Atg1 expression, has been shown to extend lifespan and improve stress tolerance in *Drosophila* [63–65]. Together, these genetic interventions emerge as a conserved genetic framework that orchestrates aging trajectories and governs health span.

In this study, we genetically modulate important aging-associated pathways mentioned above specifically in larval AMPs to understand how developmental perturbations impact long-term intestinal homeostasis. By targeting regulators of inflammaging, oxidative stress, Foxo signaling, and autophagy during development, we examine their impact on epithelial cell turnover, proliferation, lineage allocation, membrane integrity, AMP cluster organization, transcriptomic landscapes, and differentiation trajectories in the adult gut. Taken together, our work uncovers a previously unrecognized role of developmental progenitor aging in shaping long-term intestinal homeostasis and delineates the mechanistic basis of the relationship between early life stress and aging trajectories.

## Results

### Targeted genetic perturbation of molecular regulators of cellular aging in the adult midgut progenitors disrupts intestinal homeostasis

Aging is often characterized by compromised homeostatic and regenerative capabilities of stem cells [66, 67]. In this study, we elucidate the impact of genetically altering the regulators of cellular homeostasis in larval gut progenitors, known as adult midgut progenitors (AMPs), on *Drosophila* intestinal homeostasis. Previous reports suggest that increased activation of NF-κB signaling pathways driven by the overactivation of either the Toll pathway via septic injury in the fly thorax [68] or the Imd pathway through *pirk* knockdown in fly neurons [69] elevates inflammation and significantly reduces lifespan in *Drosophila* [69, 70]. Elevated Reactive Oxygen Species (ROS) levels via mitochondrial complex I knockdown have also been shown to induce oxidative stress and perturb redox homeostasis in germline stem cells of *Drosophila* testis [71]. Conversely, promoting Atg8a overexpression in aged fly brains [64] and Foxo overexpression in muscle are known to extend lifespan in *Drosophila* [72]. We employed the inflammaging approach previously reported [3, 71] to induce cellular aging in the AMPs by inducing the activation of the NF-κB pathways - Toll or Imd. Toll signaling was activated using *Toll10B*, a gain-of-function allele that constitutively activates the Toll receptor [68, 73], while Imd signaling was hyperactivated through RNAi-mediated knockdown of *Pirk*, an intracellular negative regulator of the pathway [74, 75]. In addition to inflammaging, we perturbed reactive oxygen species (ROS) homeostasis by abrogating mitochondrial complex I component, ND42, using the RNAi approach (Fig. 1A), and causing accelerated cellular aging by elevating ROS levels [76]. Conversely, cellular homeostasis was maitained by overexpressing either Atg8a, an autophagy regulator essential for autophagosome–lysosome fusion [77, 78], or Foxo, a conserved transcription factor regulating metabolism, aging, and gut integrity [79, 80] (Fig. 1A). To functionally validate the impact of these genetic perturbations on cellular homeostasis, we examined the key aging hallmarks including ROS levels, autophagy, and DNA damage, in Escargot-positive AMPs of the larval gut.

**Figure 1:**
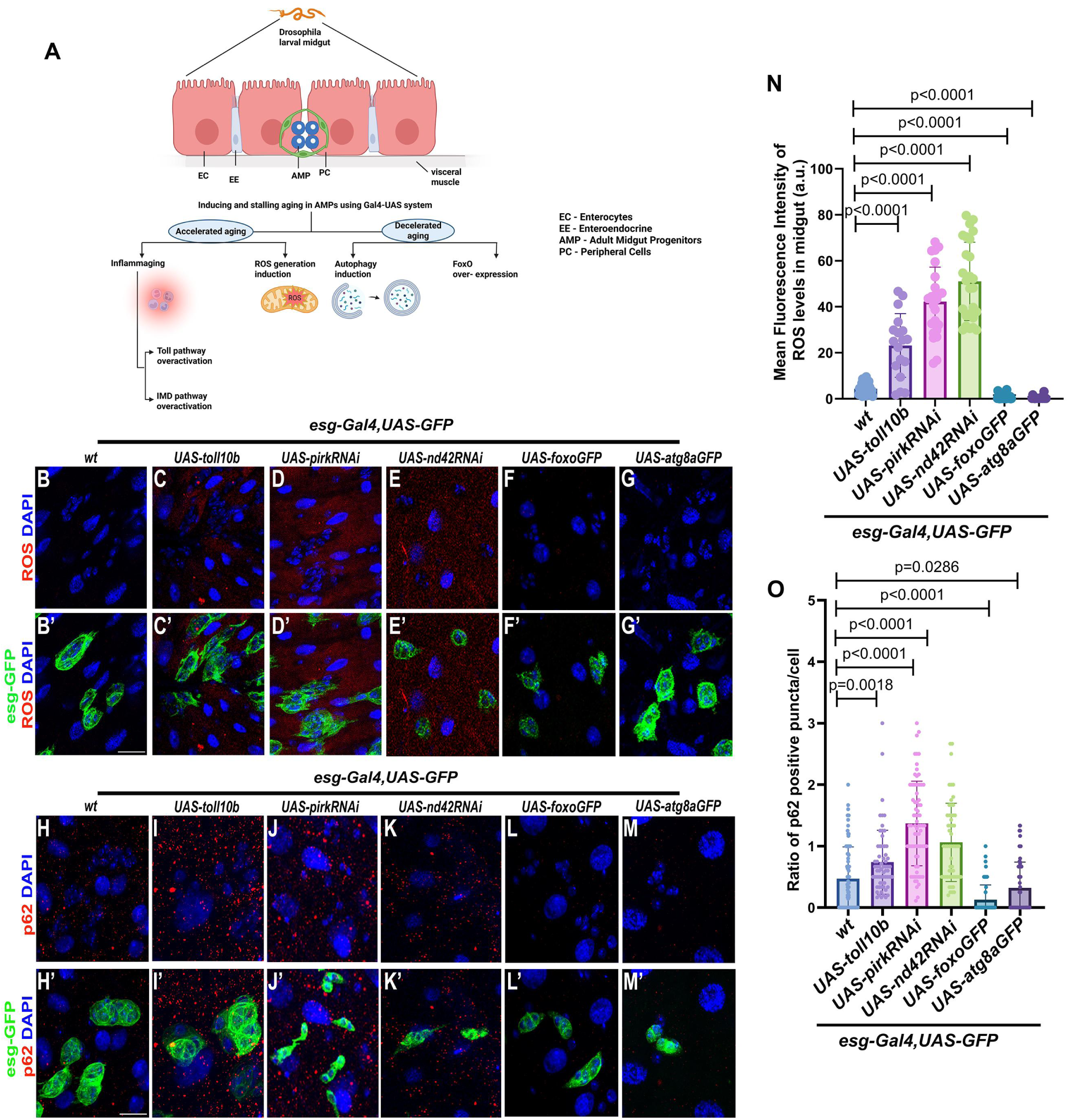
Targeted genetic perturbation of molecular regulators of cellular aging in adult midgut progenitors disrupts larval intestinal homeostasis. Schematic illustration of genetically inducing accelerated or decelerated cellular aging in larval AMPs. Cellular aging induced by over activating Toll or Imd pathways or via oxidative stress caused by ND42 knockdown. Decelerated cellular aging induced by overexpressing Foxo or Atg8a (A). ROS levels in the midgut marked by Cell-ROX Deep red (Red) upon *esg-Gal4* specific expression of *UAS-toll10b*, *UAS-pirkRNAi*, *UAS-nd42RNAi*, *UAS-foxoGFP* or *UAS-atg8aGFP* as compared to wild-type control (B-G’, N). Autophagic flux measured by quantifying the ratio of p62 positive (Red) punctae per cell upon *esg-Gal4* specific expression of *UAS-toll10b*, *UAS-pirkRNAi*, *UAS-nd42RNAi*, *UAS-foxoGFP,* or *UAS-atg8aGFP* as compared to wild-type control (H-M’, O). GFP expression (green) is driven by *esg-Gal4,UAS GFP* and nuclei stained by DAPI (blue). In ROS levels, each data point represents one ROI (N), and in p62 staining, each data point represents the average number of p62-positive puncta per cell (p/n) calculated for a single *esg* cluster. Scale bar: 5μm (B-M, B’-M’). For ROS analysis, a minimum of 6 larvae per genotype were examined and three regions of interest (ROI) were quantified per image. For p62 analysis, a minimum of 8 larvae per genotype were considered. Statistical significance was assessed using Student’s t-test with Welch’s correction. P-values are indicated in the respective graphs. Schematic (A): Created in BioRender. Khadilkar, R. (2026) https://BioRender.com/in60t75.

ROS levels, measured as mean fluorescence intensity, were significantly elevated upon Toll or Imd activation or upon ND42 knockdown and were lowered upon Foxo or Atg8a overexpression as compared to the wild-type control (Fig. 1B-G’, N). We assessed autophagy by measuring p62 levels, an adaptor protein that reflects autophagic activity due to its role in cargo delivery to Atg8-containing autophagosomes [81]. Upon genetic perturbation, the ratio of p62 positive puncta per cell was estimated, and the results indicated an accumulation of p62 positive puncta upon Toll or Imd pathway activation or upon ND42 knockdown and reduced p62 positive puncta upon over-expression of Foxo or Atg8a as compared to wild-type control (Fig. 1H-M’, O). To assess the impact of AMP-induced genetic perturbations on genomic instability, a key hallmark of aging [82], γH2AX immunostaining was performed to detect DNA damage [83–85]. Marked accumulation of γH2AX foci was observed in AMPs upon Toll or Imd pathway activation or upon ND42 knockdown, while Atg8a overexpression markedly reduced DNA damage; Foxo overexpression did not show statistically significant change compared with wild-type controls (Fig. S1A-G).

### AMP- specific perturbation of cellular homeostasis affects proliferation in the larval midgut

Several findings have shown that aging in the *Drosophila* adult gut leads to dysregulated intestinal stem cell (ISC) proliferation and differentiation, largely due to perturbations in the signaling networks that preserve ISC homeostasis [22, 30, 86, 87]. Stressors such as bacterial infection, oxidative damage, and ER stress in the *Drosophila* adult gut activate pathways like EGFR, JAK/STAT, JNK, and Hippo to drive rapid intestinal stem cell (ISC) proliferation, replacing damaged ECs and restoring epithelial integrity [86, 88–90]. Additional stressors including viral infection, hypoxia, and detergent-induced injury similarly trigger ISC hyperproliferation via Upd cytokines and TRPA1/ROS signaling, though chronic exposure risks dysplasia [89]. Although aging-associated changes in ISCs are well characterized, the impact of cellular aging or genetically induced stress on their earlier counterparts, adult midgut progenitors (AMPs), during larval development remains largely unexplored. To address this gap, the current study investigates how altering cellular aging in AMPs influences intestinal and tissue homeostasis in the *Drosophila* larval gut. AMPs are multipotent progenitors derived from embryonic endoderm and give rise to all epithelial lineages of adult midgut [25]. We initiated our analysis by genetically inducing aging specifically in AMPs, which, like ISCs, are marked by the expression of Escargot, a Snail-family transcription factor [91].

We find that constitutive activation of the Imd pathway fueled hyperproliferation marked by H3P in AMPs (Fig. 2C-C’’, G), whereas an overactivated Toll receptor had no significant effect on the AMP numbers (Fig. 2B-B’’, G) as compared to wildtype control (Fig. 2A-A’’, G). Similarly, elevated ROS levels by ND42 knockdown also promoted over proliferation (Fig. 2D-D’’, G), indicating that both chronic immune activation and oxidative stress accelerate aging-associated proliferation. Notably, the AMP-specific responses we observed are consistent with ISC behavior under similar conditions, where constitutive Imd signaling and age-associated ROS elevation have been shown to drive ISC expansion [92, 93], suggesting that inflammatory and oxidative stress pathways act as conserved proliferative cues across progenitor cell types.

**Figure 2:**
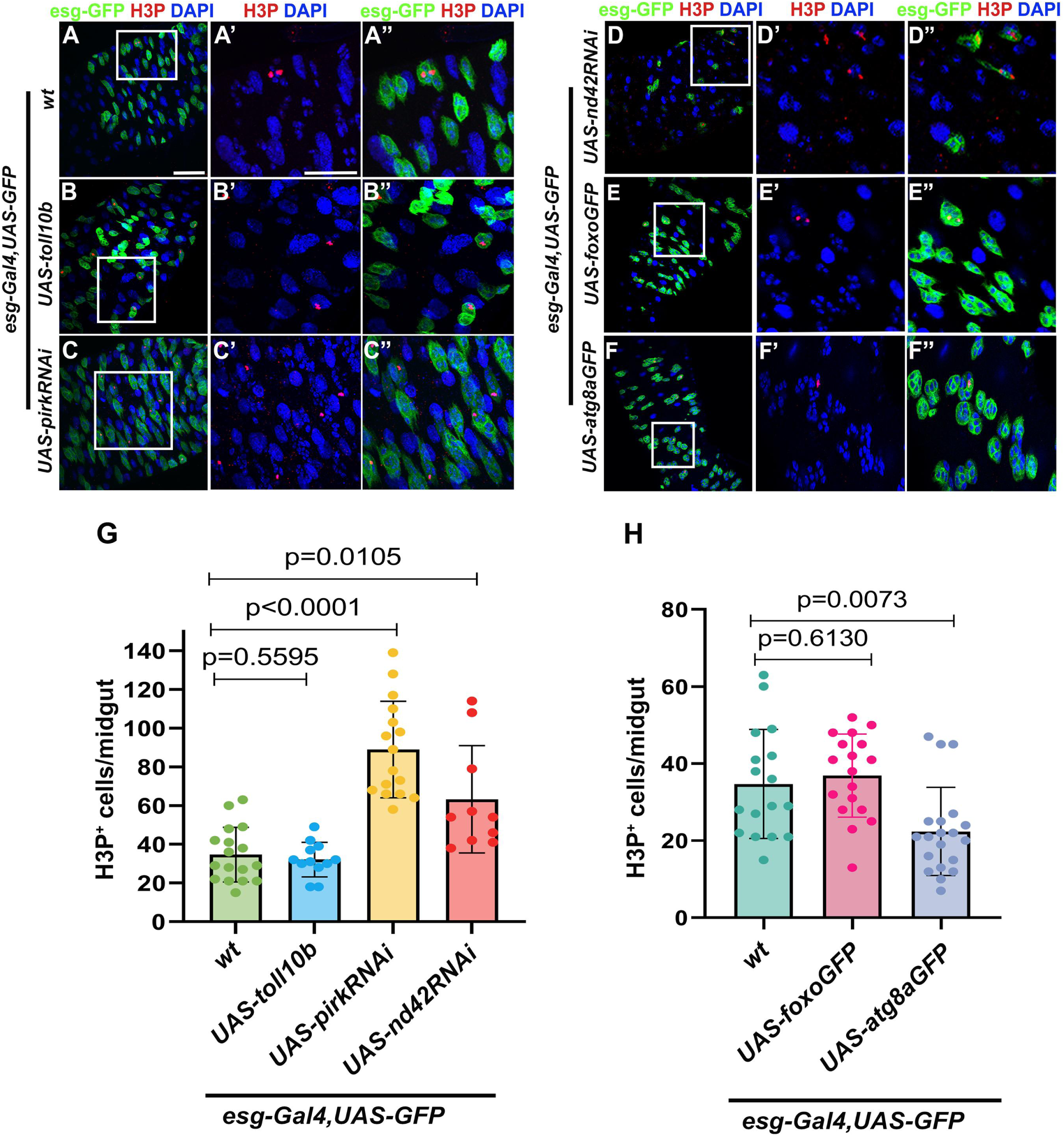
AMP- specific perturbation of cellular homeostasis affects proliferation in the larval midgut. Representative images (A–F) and quantitation (G-H) of phospho-Histone H3 (H3P, Red)-positive cells in the midgut upon *esg-Gal4* specific expression of *UAS-toll10b*, *UAS-pirkRNAi*, *UAS-nd42RNAi*, *UAS-foxoGFP,* or *UAS-atg8aGFP* as compared to wild-type control (A-F’’, G-H). Boxed regions in panels (A–F) are shown as magnified images in the corresponding panels (A’–F’, A’’-F’’). Each data point in the graph represents the number of H3P+ cells (red) per midgut. GFP expression (green) is driven by *esg-Gal4,UAS GFP* and nuclei stained by DAPI (blue). Scale bar: 20μm (A-F), 5μm (A’-F’, A’’-F’’). A minimum of 10 larvae per genotype were used for the experiment. Statistical significance was assessed using Student’s t-test with Welch’s correction. P-values are indicated in the respective graphs.

In contrast, overexpression of Atg8a significantly reduced proliferation of AMPs (Fig. 2F-F’’, H) while Foxo overexpression did not significantly alter proliferation (Fig. 2E-E’’, H) relative to the wild type (Fig. 2A-A’’, H). While previous reports stated that overexpression of Atg1 kinase in fly intestine maintains cellular homeostasis [65] and overexpression of Foxo in ISCs negatively regulates growth and proliferation in adult midgut [94], our results extend this paradigm to progenitor populations in the larval midgut during early development. Collectively, these results suggest that inflammaging and elevated ROS levels promote proliferation in both ISCs and AMPs, whereas autophagy-mediated stress responses function as conserved mechanisms to maintain gut homeostasis across developmental stages in *Drosophila*.

### Genetic modulation of cellular aging in the AMPs skews differentiation towards the enteroendocrine lineage

Previous studies have indicated that aging and stress signaling pathways can influence cellular differentiation in the intestinal epithelium. For instance, some of the studies have clearly demonstrated that aging or stress conditions alter the differentiation pattern, leading to an expansion in EE population at the expense of ECs in the adult gut [31, 32, 95, 96]. Conversely, autophagy plays a crucial role in maintaining stem cell pool [97, 98]. Overexpression of autophagy-related genes in *Drosophila* tissues, including the intestine [99] and muscles, significantly improved tissue integrity and function and systematically extends organismal lifespan [100, 101]. We therefore examined whether cellular aging-induced perturbations in AMPs impact the relative abundance of larval EEs and ECs. We observe that Toll or Imd overactivation or ND42 knockdown resulted in a non-cell autonomous increase in the number of Prospero-positive EE cells as compared to wild-type control (Fig. 3A-D’’, G). These observations parallel earlier findings showing that stress or age-related changes shift the differentiation direction towards the secretory EE lineage [31, 96]. Conversely, decelerated cellular aging, achieved by overexpressing Atg8a (Fig. 3F-F’’, H), led to a clear reduction in EE number relative to wild-type (Fig. 3A-A’’, H), whereas Foxo overexpression (Fig. 3E-E’’, H) showed a lower trend in EE differentiation but not a statistically significant change compared to wild-type control (Fig. 3A-A’’, H). These results indicate that AMP-induced genetic perturbation of molecular regulators of cellular aging can non-cell-autonomously influence surrounding midgut cells and skew the differentiation towards the EE lineage under stress conditions.

**Figure 3:**
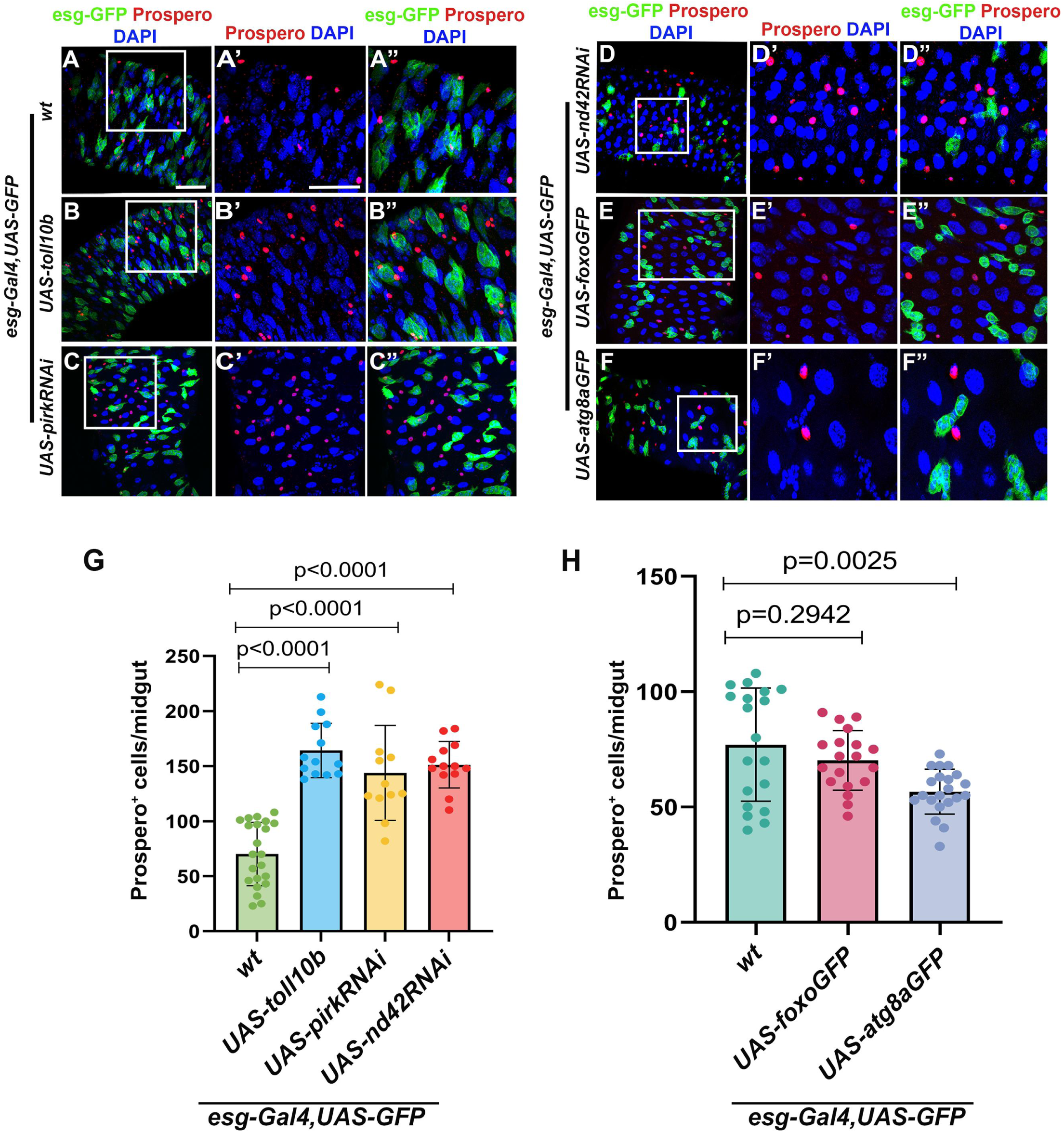
Genetic modulation of cellular aging in the AMPs skews differentiation towards the enteroendocrine lineage. Representative images (A-F) and graphical representation (G-H) of Prospero^+^ (Red) cells per midgut upon *esg-Gal4* specific expression of *UAS-toll10b*, *UAS-pirkRNAi*, *UAS-nd42RNAi*, *UAS-foxoGFP,* or *UAS-atg8aGFP* as compared to wild-type control (A-F’’, G-H). Boxed regions in panels (A–F) are shown as magnified images in the corresponding panels (A’–F’, A’’-F’’). Each data point in the graph represents the number of Prospero^+^ cells (red) per midgut. GFP expression (green) is driven by *esg-Gal4,UAS GFP* and nuclei stained by DAPI (blue). Scale bar: 20μm (A-F), 5μm (A’-F’, A’’-F’’). A minimum of 12 larvae were used for this experiment. Statistical significance was assessed using Student’s t-test with Welch’s correction. P-values are indicated in the respective graphs.

### AMP- specific genetic modulation of regulators of cellular aging controls septate junction integrity and midgut barrier function

Intact septate junction (SJ) organization in the *Drosophila* gut epithelium is crucial for maintaining barrier integrity, as it restricts paracellular leakage of solutes, microbes, and toxins into the hemolymph, thereby preventing dysbiosis, inflammation, and stem cell over proliferation. SJ disruption leads to EC detachment, polarity loss, intestinal hypertrophy, and shortened lifespan, underscoring their role in epithelial homeostasis [102, 103]. Disruption of intestinal barrier integrity during aging is an evolutionarily conserved feature across species [104, 105]. Previous findings also point out the disruption of intestinal barrier integrity in aged flies relative to young ones [34, 106]. To understand if progenitor-specific modulation of cellular aging can remodel the midgut and regulate its barrier function, we assessed the barrier integrity in the larval midgut following AMP-specific genetic interventions.

We labelled the septate junctions using Coracle, and our results demonstrate that AMP-specific Toll or Imd activation or ND42 knockdown resulted in reduced Coracle levels. In contrast, Atg8a overexpression has increased Coracle levels at the cell cortices, denoting an intact epithelial barrier as compared to the wild type, whereas Foxo overexpression did not elicit a significant change relative to the wild-type control (Fig. 4A-G). Furthermore, to functionally assess barrier integrity, the classical Smurf assay was performed by feeding larvae with a non-absorbable brilliant blue dye (Fig. 4H). We observed that the levels of Coracle correlated with the barrier integrity, wherein Toll or Imd over-activation or ND42 knockdown in the AMPs resulted in the leakage of the blue dye outside the gut, whereas the dye was limited to the gut upon AMP-specific overexpression of Atg8a or Foxo as compared to the wild type (Fig 4I). Thus, AMP-driven cellular aging disrupts barrier integrity in a manner that closely mirrors the midgut phenotypes during organismal aging in the adults.

**Figure 4:**
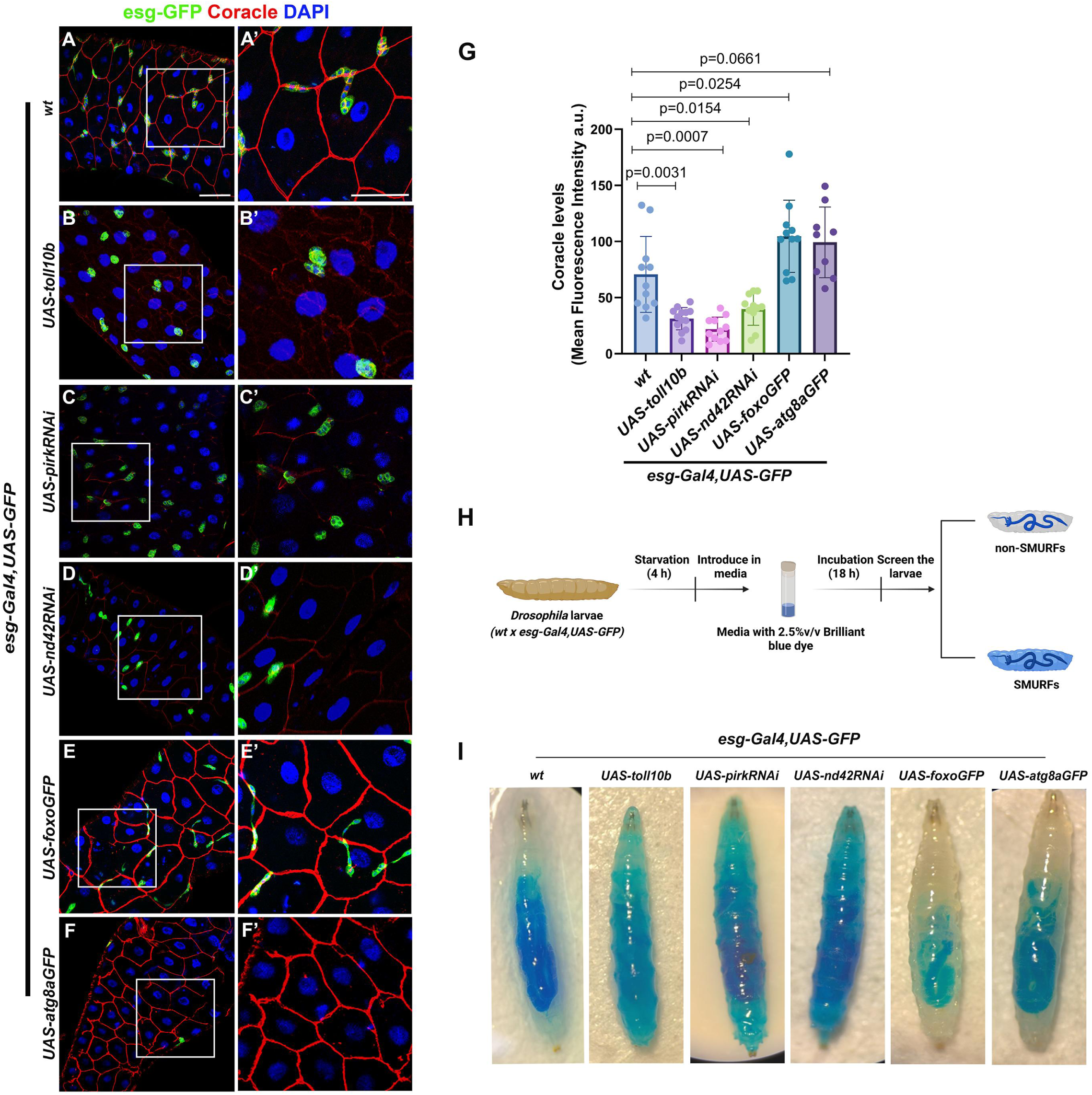
AMP- specific genetic modulation of regulators of cellular aging controls septate junction integrity and midgut barrier function. Representative images (A–F) and quantitation (G) of Coracle (Red) staining in larval midguts upon *esg-Gal4*–mediated expression of *UAS-toll10b*, *UAS-pirkRNAi*, *UAS-nd42RNAi*, *UAS-foxoGFP*, or *UAS-atg8aGFP*, compared to wild-type controls (A-F’, G). Boxed regions in panels (A–F) are shown as magnified images in panels (A’–F’). GFP (green) expression is driven by *esg-Gal4,UAS-GFP*, and nuclei stained by DAPI (blue). Schematic of the Smurf assay in larvae (H). Representative Smurf assay images from larvae with *esg-Gal4,UAS-GFP* mediated expression of *UAS-toll10b*, *UAS-pirkRNAi*, *UAS-nd42RNAi*, *UAS-foxoGFP,* or *UAS-atg8aGFP* as compared to wild-type controls (I). Scale bar: 20μm (A-F), 5μm (A’-F’). A minimum of 8 larvae were considered for Coracle quantitation. Each data point represents the mean fluorescence intensity calculated from 5 individual measurements per image. For the SMURF assay, a minimum of 8 larvae were considered. Statistical significance was determined using Student’s *t*-test with Welch’s correction. P-values are indicated in the graph. Schematic (H): Created in BioRender. Khadilkar, R. (2026) https://BioRender.com/sn1ko47

To independently validate the genetic perturbations, we next employed chemical interventions using paraquat and rapamycin and checked Coracle levels (Fig. S2A). Paraquat induces oxidative stress that mimics aging phenotypes and accelerates cellular damage in *Drosophila* tissues and organs. Rapamycin, an mTOR inhibitor, promotes autophagy and extends lifespan, serving as an anti-aging intervention to counteract stress-induced defects [107–109]. Midguts from paraquat-fed larvae showed a reduction in Coracle levels, whereas Rapamycin-fed larvae displayed elevated Coracle levels in the midgut as compared to the midguts from the vehicle-treated larvae (Fig. S2B-H). These findings are consistent with previous reports demonstrating that paraquat disrupts epithelial repair [110], whereas rapamycin promotes intestinal barrier repair [111].

We then performed the Smurf assay on the larvae fed with Paraquat or Rapamycin, and found that paraquat-fed larvae showed a leaky gut phenotype while Rapamycin-fed larvae had their gut barrier intact as compared to the control-treated larvae (Fig. S2G-I). Collectively, these observations are consistent with previous literature, which states that age-related stem cell dysfunction compromises barrier integrity [104, 112] while autophagy plays a protective role in epithelial barrier homeostasis [113]. These findings align with our speculations that early progenitor-specific genetic interventions can profoundly impact intestinal homeostasis.

### Chemical-based modulation of aging affects larval intestinal homeostasis

Since AMP-specific genetic perturbation showed increased DNA damage, increased AMP proliferation, and altered lineage allocation, we next sought to validate these findings by modulating aging pathways using chemical interventions. Paraquat or Rapamycin administration was done at the organismal level to chemically modulate cellular aging in the *Drosophila* larvae. Previous literature has established that Paraquat, a mitochondrial toxicant, induces oxidative stress [114, 115], causes DNA damage [116] and its exposure substantially induces ROS production and suppresses activation of the longevity factor Foxo3, in cultured cardiomyocytes and cardiac tissue [117].

Paraquat or Rapamycin was fed to the larvae for 12 hours after a 2-hour starvation window, and midguts were analyzed (Fig. 5A). Paraquat administration to the larvae led to an increase in H3P-positive mitotically active cells and an increase in Prospero-positive EE population in the midguts relative to its vehicle control (Fig. 5B-G), indicating that Paraquat-mediated ROS production perturbs intestinal homeostasis. Given that oxidative stress is a major driver of genomic instability, we next examined DNA damage under these conditions (Fig. S3A). Midguts in paraquat-fed larvae showed elevated DNA damage marked by increased γH2AX-positive Escargot-GFP cells compared to the vehicle-treated control (Fig. S3B-C’’, F). In contrast, Rapamycin treatment resulted in a decrease in H3P-positive mitotically active cells and Prospero-positive EEs as compared to the vehicle-treated control (Fig. 5H-M). Also, rapamycin-fed larvae displayed a decrease in γH2AX-positive Escargot-GFP cells as compared to the vehicle control (Fig. S3D-G).

**Figure 5:**
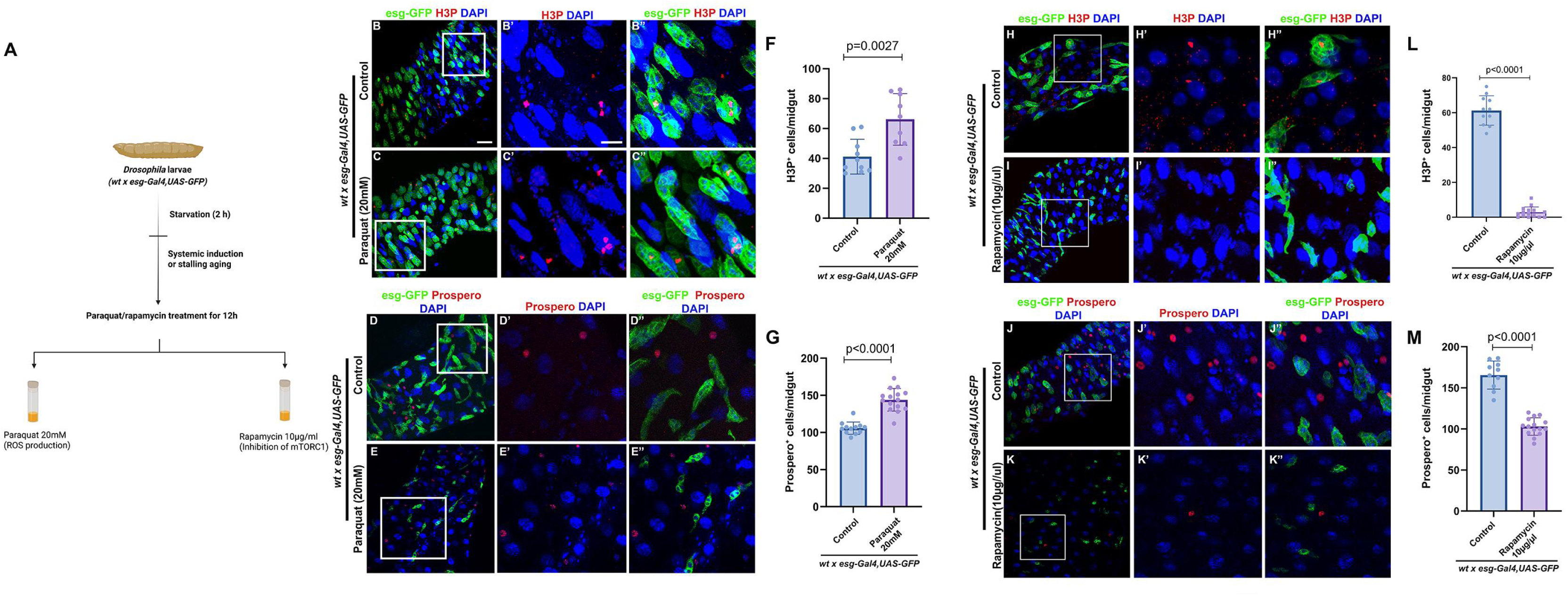
Chemical-based modulation of aging affects larval intestinal homeostasis. Schematic illustration of chemical treatment of the larvae (A). Representative images (B-C, H-I) and quantitation of cells stained with phosphorylated histone H3 (H3P^+^, Red) in midgut upon Paraquat (20mM) (B-C’’, F) or Rapamycin (10μg/ml) treatment (H-I’’, L), respectively. Representative images showing Prospero^+^ enteroendocrine cells (D–E’’, J–K’’) and their quantitation (F, L) upon Paraquat (D–E’’) or Rapamycin (J–K’’) treatment. Boxed regions in panels (B-E, H-K) are shown as magnified images in the corresponding panels (B’–E’’, H’-K’’). Each data point in the graphs (F-G, L-M) represents H3P^+^ or Prospero^+^ cells (Red) per midgut. For quantitation of H3P^+^ or Prospero^+^ cells, Paraquat and Rapamycin treatment conditions were analyzed as compared to the vehicle control (F, G, L, and M). GFP (green) expression is driven by *esg-Gal4,UAS GFP,* and nuclei stained with DAPI (blue). Scale bar: 20μm (B-E, H-K), 5μm (B’-E’, B’’-E”, H’-K’, H”-K”). For H3P and Prospero staining, a minimum of 10 larvae were analyzed for Rapamycin treatment and 9 larvae for the Paraquat experiment. Statistical significance was assessed using Student’s t-test with Welch’s correction. P-values are indicated in the respective graphs. Schematic (A): Created in BioRender. Khadilkar, R. (2026) https://BioRender.com/iw99214.

Collectively, these results demonstrate that chemical modulation of the cellular aging systemically affects larval intestinal homeostasis phenocopying the outcomes observed upon AMP-specific genetic perturbations.

### Transcriptomic profiling of larval midguts with AMP-specific genetic interventions recapitulates molecular hallmarks of aging

To investigate the underlying molecular mechanisms that are impacted upon AMP-specific genetic modulation of cellular aging, we performed mRNA sequencing of *Drosophila* larval midguts subjected to AMP- specific Toll or Imd over-activation, ND42 knockdown, or Foxo or Atg8a overexpression (Fig. 6A). Principal-component analysis (PCA) results revealed a separation between aging (Toll overexpression, *Pirk* and ND42 knockdown) and anti-aging genotypes (Foxo and Atg8a overexpression), with overall genotype specific clustering; however moderate heterogeneity was observed among aging replicates, with one Pirk and Toll replicate clustering closer to the Foxo group (Fig. 6B, Fig. S4A and S5A). This divergence demonstrated that aging and anti-aging interventions in the AMPs induce significant changes in gene enrichment profile in the larval midgut. To further identify genes differentially regulated between aging and anti-aging conditions, differential gene expression (DEG) analysis was performed comparing the aging genotypes with the anti-aging genotype (Foxo overexpression), and the results were visualized using volcano plots. At a significance threshold of p < 0.05, 189 genes were identified upon Toll over-activation versus Foxo overexpression (Fig. 6C), 116 genes upon *pirk* knockdown (Fig. S4B), and 280 genes in ND42 knockdown (Fig. S5B) using Foxo as a reference genotype.

**Figure 6:**
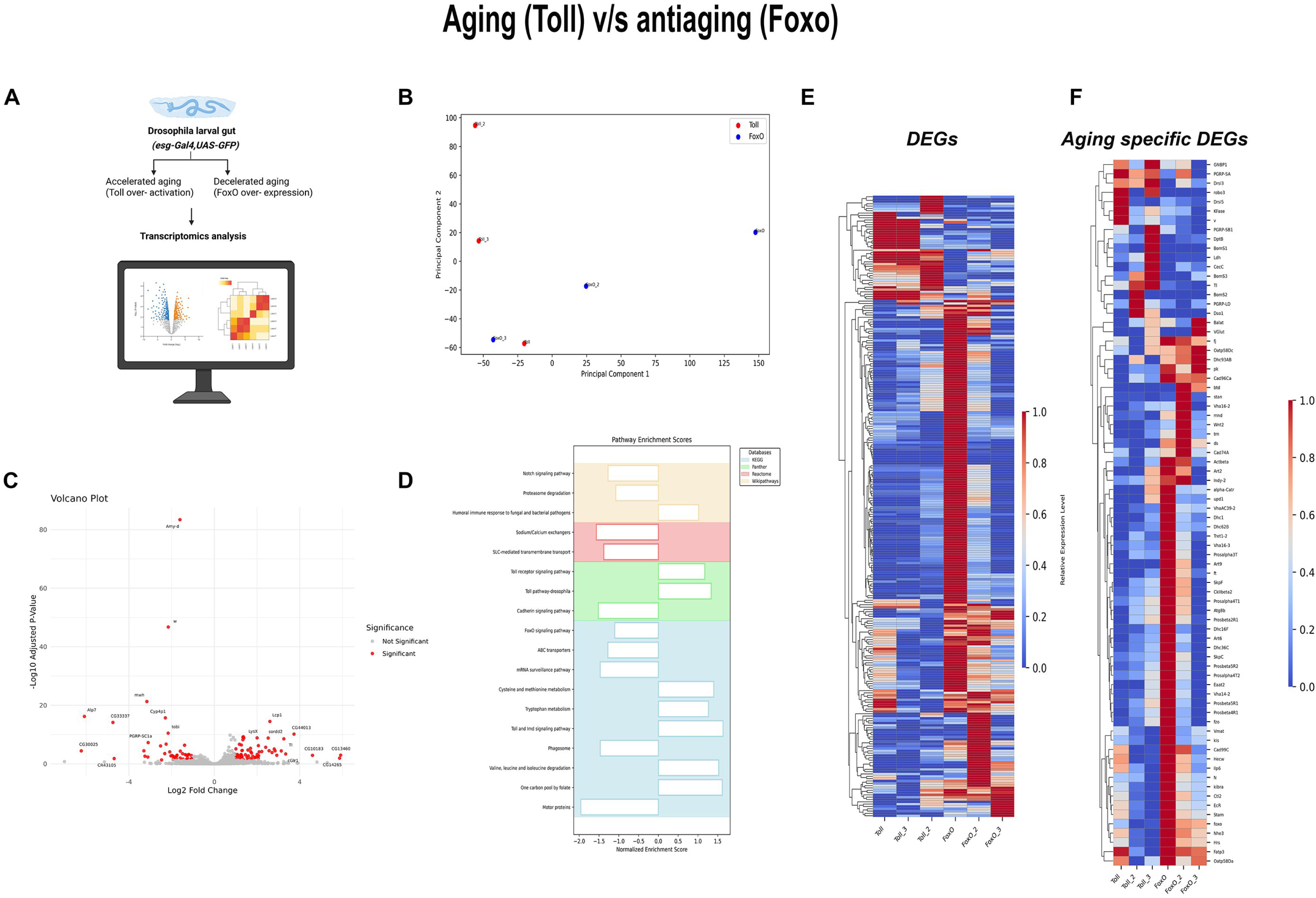
Differential gene expression and pathway enrichment analyses highlight aging-associated transcriptional changes in AMP-specific Toll activation versus Foxo overexpressing midguts. Schematic representation of experimental flow of bulk RNA-seq analysis using *Drosophila* larval gut (A). Principal Component Analysis (PCA) plot for biological replicates of Toll receptor-activated (test), and Foxo overexpressed (reference) genotype gut cells (B). Volcano plot representing significantly deregulated genes (C). Barplot representing pathway enrichment scores obtained from Gene Set Enrichment Analysis performed with KEGG, Panther, Reactome, and Wikipathways database (D). Heatmap representing differentially expressed genes involved in hallmark aging pathways (E). Heatmap representing overall differentially expressed genes (F). Schematic (A): Created in BioRender. Khadilkar, R. (2026) https://BioRender.com/728ja2d.

Gene set enrichment analysis (GSEA) of aging genotypes (Toll overactivation, *Pirk* and ND42 knockdown), using the anti-aging genotype (Foxo overexpression) as a reference, revealed a pronounced upregulation of genes associated with the humoral immune response as well as Toll (Tl, Drsl3, BomS1) and Imd signaling pathways (PGRP-SB1, CecC, DptB) which are characteristic of classical aging phenotypes (Fig. 6C, Fig. S4B, S5B). Consistent with earlier reports, immune signaling pathways are known to remain chronically activated under aging conditions [40, 118]. In addition, and in agreement with previous studies [119–121], significant dysregulation of several metabolic processes including valine, leucine, and isoleucine metabolism, as well as methionine and cysteine metabolism was observed in aging genotypes compared with anti-aging ones (Fig. 6D, Fig. S4C, Fig. S5C). Notch signaling was markedly downregulated in aging genotypes (Fig. 6D, Fig. S4C, Fig. S5C). Although primarily characterized in adult guts [122, 123], our results suggest a similar role for Notch in the larval midgut, where diminished Notch activity likely contributes to the increased EE differentiation observed in aging conditions. In line with previous findings [124–126], proteasome-mediated degradation (Fig. 6D; Fig. S5C) and protein ubiquitination (Fig. S4C) were also downregulated in aging genotypes, indicating defective proteostasis. Additionally, aging genotypes displayed downregulation of ABC transporters, sodium/calcium exchangers, SLC-mediated membrane transport, cadherin signaling and mRNA surveillance pathways (Fig. 6D; Fig. S4C; Fig. S5C), reflecting widespread impairment in cellular transport, adhesion, and overall epithelial integrity.

To obtain a comprehensive overview of global transcriptional changes across the different genetic conditions, heatmaps were generated using DESeq2-normalized counts. The heatmap analysis revealed upregulation of innate immune response genes and downregulation of genes involved in gut integrity and barrier function, proteostasis, cytoskeletal organization, ion transport, and developmental signaling pathways in the Toll overexpression genotype (Fig. 6E, F). *Pirk* knockdown condition showed upregulation of origin recognition complex (ORC) and minichromosome maintenance (MCM) genes, indicating damage-induced compensatory replication initiation, while compromising CPSF and mRNA processing fidelity [127, 128] (Fig. S4D, E). In ND42 knockdown, glutathione S-transferase (GST) genes were significantly upregulated (Fig. S5D, E), indicating enhanced antioxidant response, consistent with earlier reports [129, 130]. Conversely, key gene sets involved in proteasomal degradation [131], cytoskeletal organization [132], barrier integrity [133], and lysosomal and autophagy pathways [134] were downregulated in both *pirk* (Fig. S4D, E) and ND42 knockdown (Fig S5D, E) conditions. All aging genotypes were compared relative to Foxo overexpression for the GSEA analysis. Collectively, these transcriptional changes indicate that genetically induced stress in larval AMPs recapitulates key molecular signatures of intestinal aging, providing mechanistic insight into how stress-induced aging pathways in progenitors influence gut homeostasis.

### Genetic modulation of cellular aging in the AMPs alters the structure and spatial arrangement of AMP clusters

In line with our earlier observations that AMP-specific modulation of cellular aging leads to a non–cell-autonomous increase in EE cell numbers under accelerated aging and a reduction under decelerated aging, we next investigated whether aging also affects AMP cluster number and morphology (Fig. 7A). Under accelerated aging conditions, the area occupied by AMP clusters was significantly reduced in aging genotypes, except the Toll over-activation conditions, which did not show a significant change, whereas Foxo over-expression exhibited cluster areas comparable to wild-type controls and Atg8 over-expression showed a decrease in area occupied by AMP clusters (Fig. 7H). Consistent with these observations, the total number of AMP clusters per midgut was also significantly reduced under accelerated aging, whereas decelerated aging did not show a significant increase in cluster number (Fig. 7B-G’, I).

**Figure 7:**
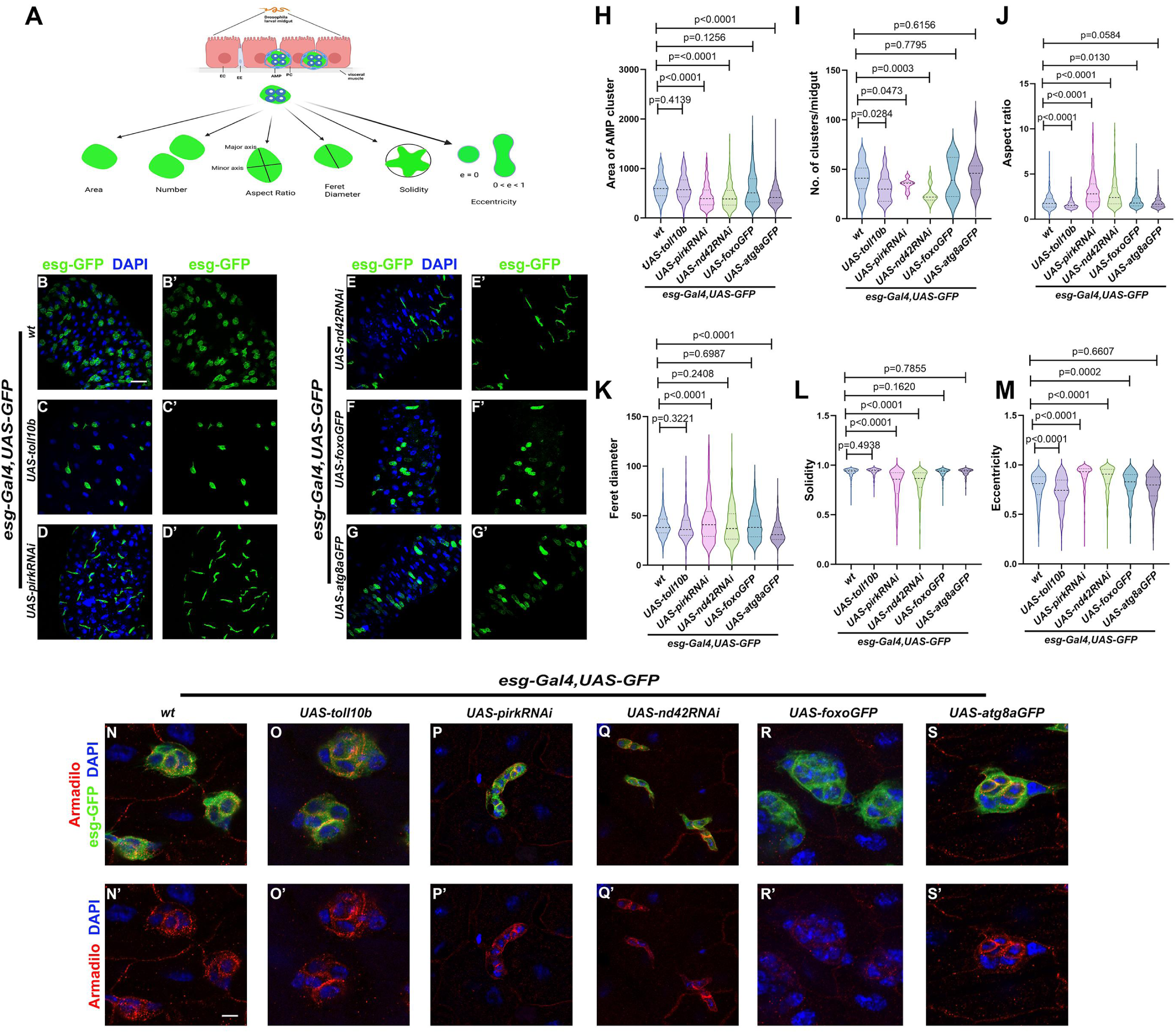
Genetic modulation of cellular aging in the AMPs alters the structure and spatial arrangement of AMP clusters. Schematic overview of the quantitative morphometric analysis. The top illustration depicts the *Drosophila* larval posterior midgut epithelium, with adult midgut progenitors (AMPs) and peripheral cells (PCs) highlighted. The lower part depicts the key morphometric parameters that were quantified (A). Representative images of midguts (B-G’). Progenitor cells (AMPs) are marked by GFP expression under the control of an escargot driver (*esg-Gal4,UAS-GFP*). Escargot-GFP (Green) positive AMP clusters upon *escargot-Gal4, UAS-GFP* specific expression of *UAS-toll10b* (C, C’), *UAS-pirkRNAi* (D, D’), *UAS-nd42RNAi* (E, E’), *UAS-foxoGFP* (F, F’) or *UAS-atg8aGFP* (G, G’) as compared to the wild-type control (B-B’). Quantitation of AMP cluster numbers and morphometric parameters for the genotypes shown in (B-G’). Violin plots illustrate the distribution for each measured parameter: Area of AMP cluster (H), Number of cell clusters per midgut (I), Aspect Ratio (J), Feret Diameter (K), Solidity (L), and Eccentricity (M). Representative images of AMPs stained for Armadillo (red) to visualize AMP shape (N-S) and cellular organization upon *esg-Gal4,UAS-GFP* expression of *UAS-toll10b* (O, O’), *UAS-pirkRNAi* (P, P’), *UAS-nd42RNAi* (Q, Q’), *UAS-foxoGFP* (R, R’) or *UAS-atg8aGFP* (S, S’) relative to the wild-type control (N, N’). Armadillo and DAPI channel from the same regions, emphasizing overall cell shape (N’-S’). GFP (green) expression is driven by *esg-Gal4,UAS-GFP,* and nuclei stained by DAPI (blue). A minimum of 12 larvae were used for each parameter. Scale bar: 20μm (B-G, B’-G’) 5μm (N-S, N’-S’). Statistical significance was assessed using Student’s t-test with Welch’s correction. P-values are as indicated in the graphs. Schematic (A): Created in BioRender. Khadilkar, R. (2026) https://BioRender.com/ky80j0j.

Morphometric analyses revealed that accelerated aging led to an increase in the aspect ratio (Fig. 7J) and feret diameter of AMP clusters (Fig. 7K), indicating elongation and expansion in cluster shape (Fig. 7D-D’, E-E’). Conversely, decelerated aging caused a decrease in aspect ratio (Fig. 7J), suggesting more compact clusters (Fig. 7B-B’, C-C’, F-F’, G-G’). Additionally, solidity decreased (Fig. 7L) and eccentricity of clusters increased with accelerated aging (Fig. 7M) whereas eccentricity increased in Foxo overexpression and showed non-significance in Atg8a overexpression conditions (Fig. 7M), reflecting changes in cluster compactness and shape regularity.

To determine whether age-dependent alterations in cluster morphology are reflected in individual progenitor cells, we stained larval midguts for Armadillo, a junctional marker that delineates cell boundaries and enables visualization of AMP cell shape (Fig. 7N-S’). In wild-type controls (Fig. 7N, N’), AMPs exhibited well-defined cell boundaries and a compact, orderly morphology. In contrast, accelerated cellular aging resulted in irregular AMP shapes and disordered cellular organization (Fig. 7P-Q’); though, Toll10b overactivation (Fig. 7O-O’) did not result in detectable irregularities in AMP morphology. Notably, decelerated aging (Fig. 7R-S’) maintained a control AMP-like morphology with well-defined cell boundaries and coherent cellular organization as compared to the wild-type control (Fig. 7N-N’). These results suggest that perturbation of cellular aging not only modulates the number but also the morphology of AMP clusters/islets, which may underlie altered differentiation dynamics and turnover in the adult midgut during aging.

Intriguingly, age-associated changes in AMP cluster morphology have not been previously reported. However, deregulated progenitor clustering has been described in the aging *Drosophila* midgut, where aberrant clustering of EC progenitors has been identified as an early-life biomarker of aging and is associated with impaired progenitor identity and differentiation [135]. In line with this, our data show that genetic modulation of cellular aging programs significantly alters both the number and morphology of larval AMP clusters, suggesting that aging influences the spatial organization of progenitors at early developmental stages.

### Early life AMP-specific cellular aging-associated stress exerts long-term effects on adult intestinal homeostasis

Given that AMP-specific genetic perturbations and systemic chemical intervention elicited aging-like phenotypes in the midgut in larval stages, we next examined whether early life AMP-specific cellular aging affects intestinal homeostasis in adulthood. To address this, we induced AMP-specific genetic perturbations starting from early development and subsequently analyzed adult midguts at 7- or 30 days post eclosion (DPE).

All previously described genotypes successfully eclosed into adult flies, except for AMP-specific ND42 knockdown, which failed to eclose and was therefore excluded from adult-stage analysis. All the flies were analyzed at 7 days post eclosion (DPE), and 30 DPE to assess age-dependent intestinal differentiation trajectories (Fig. 8A). At 7 DPE, constitutive activation of the Toll or Imd pathways resulted in a pronounced shift in intestinal differentiation towards the EE lineage (Fig. 8C-D’, G). In contrast, Foxo or Atg8a overexpression-maintained EE proportions (Fig. 8E-F’, G) comparable to the wild-type controls (Fig. 8B-B’, G). At 30 DPE, a similar pattern was observed with elevated EE numbers in aging-associated genotypes (Fig. 8I-J’, M) relative to anti-aging genotypes (Fig.8K-L’, M) and the wild-type control (Fig. 8H-H’, M).

**Figure 8:**
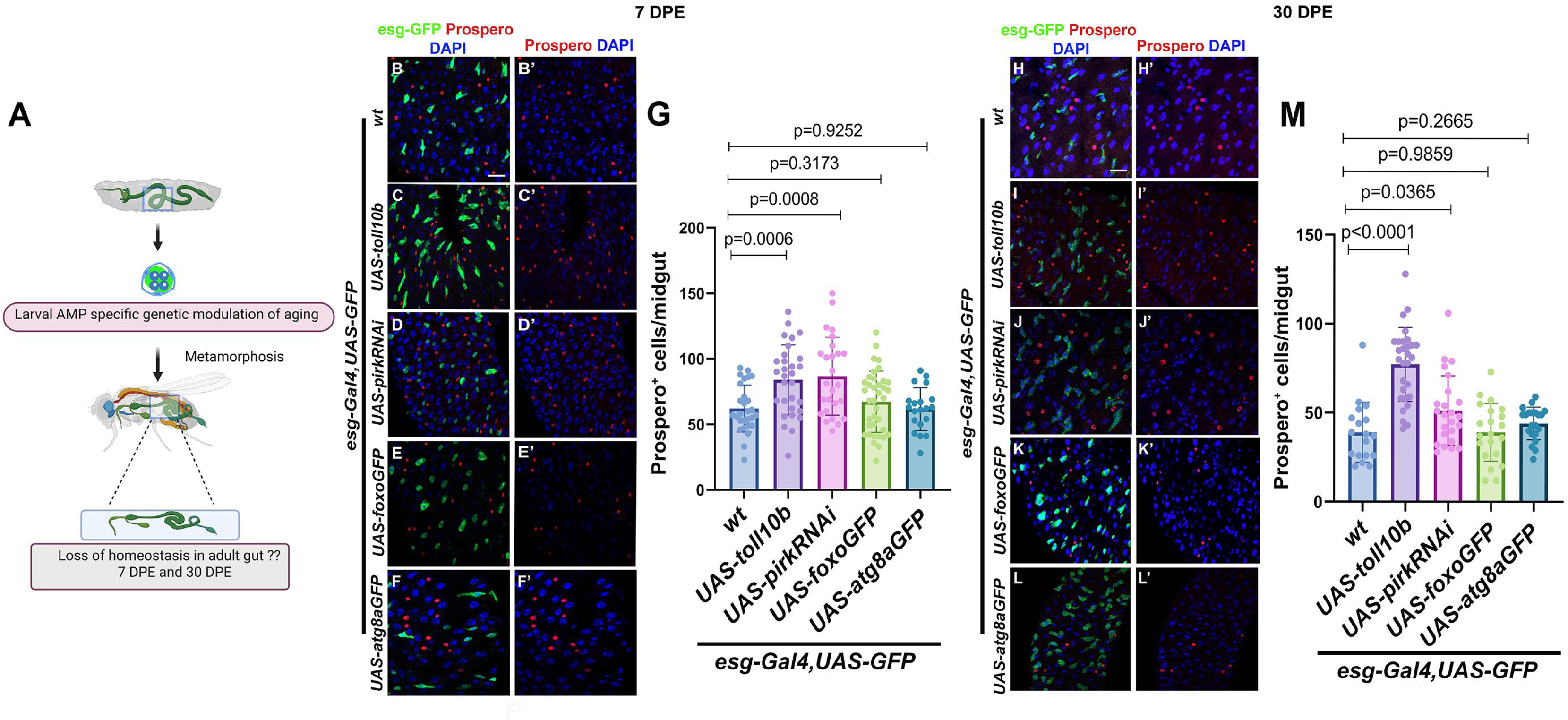
Early life AMP-specific cellular aging-associated stress exerts long-term effects on adult intestinal homeostasis. Schematic illustrating the impact of AMP-specific genetic modulation on adult gut homeostasis (A). Representative images of Prospero^+^ enteroendocrine cells at 7 days post-eclosion (DPE) and 30 DPE in adult midgut with *esg-Gal4,UAS-GFP* driven expression of *UAS-toll10b* (C–C’, I–I’), *UAS-pirkRNAi* (D–D’, J–J’), *UAS-foxoGFP* (E–E’, K–K’), or *UAS-atg8aGFP* (F–F’, L–L’) as compared to wild-type control (B–B’, H–H’). Graphs show quantitation of Prospero^+^ cells per midgut at 7 DPE (G) and 30 DPE (M). Each data point in the graph indicates the no. of Prospero^+^ cells quantified in the half midgut region. GFP (green) expression is driven by *esg-Gal4,UAS-GFP,* and nuclei stained by DAPI (blue). Scale bar: 20μm (A-L, A’-L’). A minimum of 10 adults were used for 7 and 30 DPE experiments. Statistical significance was assessed using Student’s t-test with Welch’s correction. P-values are as indicated in the graphs. Schematic (A): Created in BioRender. Khadilkar, R. (2026) https://BioRender.com/4ozzemw.

In addition, we also quantified ECs numbers in adult midguts to assess whether the observed differentiation shifts correspond with reduced EC abundance. While aging-associated genotypes displayed a non-significant decrease in EC numbers (Fig. S6C-D, G), anti-aging genotypes exhibited a significant increase in EC abundance (Fig. S6E-F, G) as compared to the wild-type (Fig. SB, G). These observations are consistent with earlier reports wherein age-associated changes in *Drosophila* intestine caused dysregulated stem cell differentiation, accompanied by altered lineage trajectories towards EE fate [31, 96, 136].

Taken together, our findings show that AMP-specific cellular aging perturbations impact intestinal homeostasis during development and persist to cause age-dependent defects in adult gut homeostasis following metamorphosis (Fig. 9). Our study provides important insights into how cellular aging at the level of stem/progenitor cells can regulate overall organ homeostasis and its response to stress. This has useful implications in understanding how stress at early developmental stages could transition into a disease scenario at a later stage in life.

**Figure 9:**
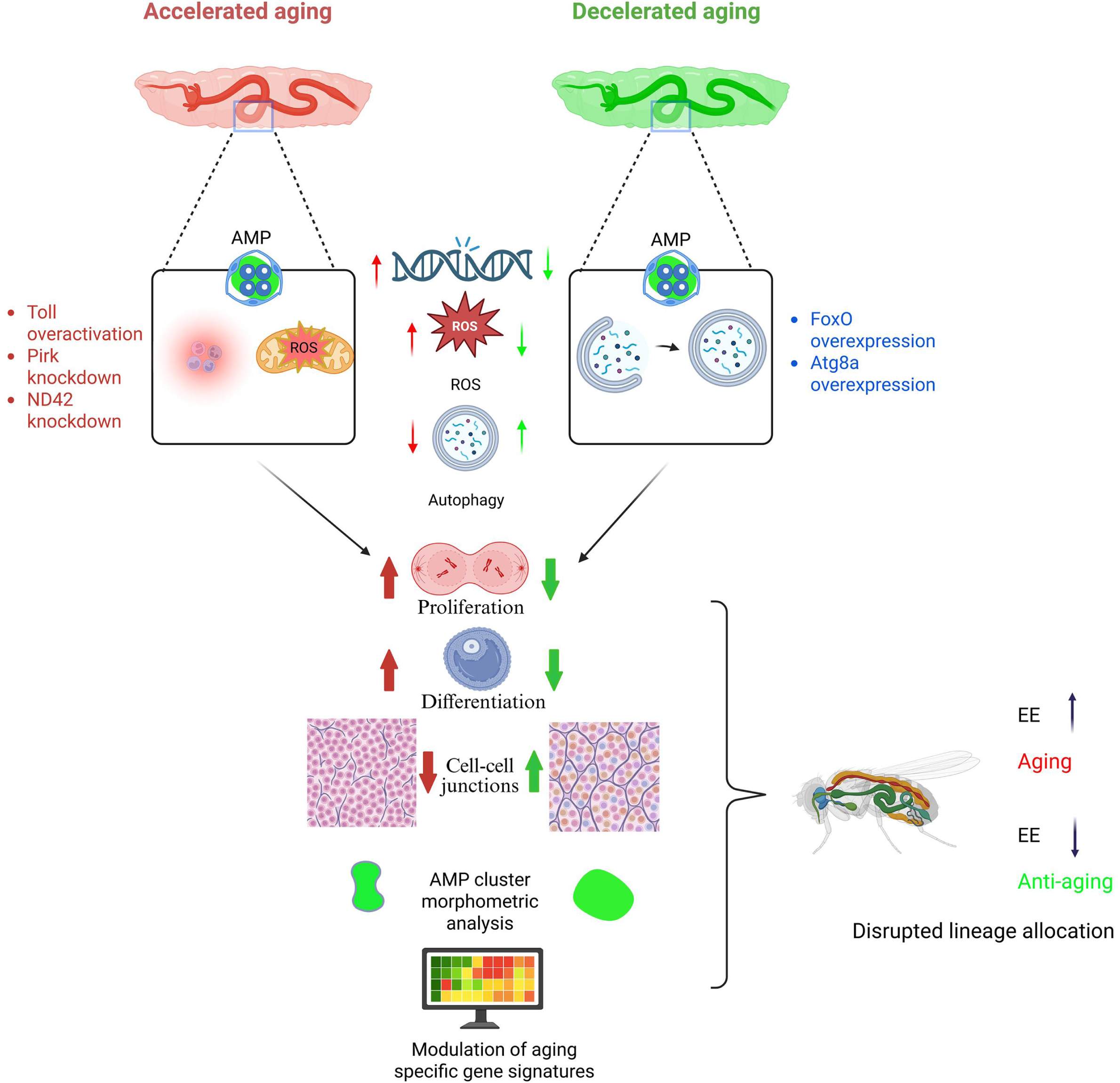
Developmental programming of cellular aging determines adult gut homeostasis. Our mechanistic model depicts how genetic acceleration of cellular aging in larval AMPs elevates oxidative stress, impairs autophagy, disrupts epithelial architecture and lineage specification, alters AMP islet morphology, and modulates aging specific gene signatures, ultimately driving adult EE lineage bias and epithelial dysfunction. Created in BioRender. Khadilkar, R. (2026) https://BioRender.com/n7hu0c0.

## Discussion

Aging is a time-dependent pathophysiological process in which stem-cell exhaustion emerges as a major hallmark, marked by compromised self-renewal and aberrant differentiation across tissues, including the intestinal epithelium [137, 138]. While aging-related decline has been extensively studied in adult intestinal stem cells (ISCs) [1, 30, 139–142], whether aging-associated stress during earlier development impacts long-term intestinal homeostasis has remained largely uncharacterized. Here, we leverage the *Drosophila* intestine as a model and elucidate that genetic modulation of cellular aging in AMPs markedly impacts intestinal homeostasis during development and later in adulthood.

AMPs give rise to all epithelial lineages of the adult intestine [26, 143]; yet, how aging-associated stress shapes AMP biology, including their proliferation, non-cell autonomous effects on other cellular subsets, membrane integrity, transcriptional changes, and ultimately adult gut function, remains largely undetermined. Our observations demonstrate that AMPs elicit core features of accelerated aging in response to chronic immune activation or upon ND42 knockdown including elevated oxidative stress, impaired autophagy, genomic instability, and disrupted proliferative control. Notably, while constitutive activation of the Imd pathway [47, 86] and elevated ROS levels due to mitochondrial complex I knockdown [89] significantly increased AMP proliferation, Toll pathway overactivation did not elicit a comparable proliferative response. Likewise, although Foxo overexpression effectively preserved cellular homeostasis by maintaining redox balance and autophagic activity, it does not suppress AMP proliferation under basal conditions. AMP-specific Foxo over-expressing midguts may respond to stress or insult differently, giving them an advantage that needs to be explored. These findings indicate that distinct aging-associated pathways differentially regulate proliferative outputs in AMPs, and that immune or longevity signals do not uniformly influence progenitor cell cycling.

We show that stress in AMPs acts non-cell-autonomously to drive EE expansion in the larval gut. Genetic triggers of inflammation (Toll/Imd activation) or mitochondrial dysfunction (*ND42* knockdown) both resulted in EE cells expansion. These results mirror observations in aging adults, where stress-induced lineage shifts favor secretory cells [31, 32, 122, 123, 144]. Given that diminished Notch signaling typically mediates this bias in adult progenitors [30, 31, 145–148] it likely plays a similar role in the larval response. Notably, the insignificant effect of *Foxo* overexpression on EE numbers suggests that Foxo’s influence on differentiation is highly context-specific [145] and may be less potent during larval stages than in later life. Systemic chemical modulation of aging pathways independently validated these findings. Paraquat-induced oxidative stress [149] recapitulated genetically accelerated aging, causing elevated AMP proliferation, DNA damage, and altered lineage allocation, consistent with ROS as a key mediator of aging [150] in the gut. In contrast, rapamycin-mediated inhibition of mTOR resulted in decreased AMP proliferation, suppressed EE numbers, and reduced DNA damage, closely reflecting the protective effects of Foxo or Atg8a overexpression [30, 151, 152]. Collectively, these findings reinforce that oxidative stress and mTOR-dependent signaling are key regulators of AMP proliferation and lineage balance during development.

Aging-associated stress was also accompanied by disrupted septate junction organization and compromised epithelial barrier integrity. Toll or Imd pathway activation and ND42 knockdown resulted in Coracle down-regulation, indicating that modulation of cellular aging programs can alter epithelial architecture during development. Decreased Coracle levels also correlated with compromised barrier integrity. Future investigation would be required to understand how genetic modulation of cellular aging in AMPs can have an impact on overall septate junction organization and barrier function in the intestine. Consistent with these observations, age-associated epithelial barrier dysfunction is a characteristic hallmark of aging and is strongly coupled with inflammatory signaling and progressive tissue decline [34, 133, 141, 153]. Our findings demonstrate that similar epithelial defects can emerge much earlier during larval midgut development, suggesting that aging-associated epithelial defects may be developmentally programmed rather than solely acquired later in life.

Bulk transcriptomic profiling of larval midguts revealed that accelerated aging in AMPs induces broad gut remodeling by showing typical aging signatures like metabolic imbalance [139], impaired proteostasis [154, 155], altered cytoskeletal organization [132, 156], compromised barrier integrity [157], and activation of developmental signaling [31], particularly under conditions of NF-KappaB signaling or ROS induction. Aging genotypes showed significant alterations in molecules important in maintaining proteostasis, for example, diminished expression of key ubiquitin-proteasome system molecules, including proteasome subunits and E3-ligase associated factors such as Prosbeta2R1, Prosalpha3T, SkpF, indicative of impaired protein turnover, and quality control. In parallel, various metabolic pathways, including molecules in amino acid metabolism (mnd, VGlut, Art6), were downregulated, indicating that early progenitor aging initiates broad remodeling of gut physiology akin to adult aging [2]. Aging genotypes also exhibited compromised mRNA surveillance pathways [158], potentially exacerbating proteotoxic stress and differentiation defects. Consistent with impaired proteasome activity, reduced expression of mitophagy and autophagy-related genes (e.g., Atg8b) reflects failure in turnover of damaged proteins and mitochondria.

Consistent downregulation of Notch signaling (N, Dl) in aging genotypes suggests that age-related signaling cascades can be developmentally programmed early in life, persist to impact adult-tissue homeostasis, and skews progenitor differentiation toward the EE lineage [123, 159]. Aging genotypes also exhibited reduced expression of cytoskeletal organization genes (Dhc36C, Dhc62B, Dhc1, robl37BC, robl22E), indicating compromised structural integrity, which may impair progenitor cell shape, cell-cell adhesion, and epithelial architecture. Genes governing epithelial membrane architecture (Kibra, Starrynight, Cad99Ca) and progenitor identity (kismet and buttonhead), crucial for maintaining gut epithelial architecture and stress resilience [155, 157], were also downregulated, providing mechanistic insight into the septate junction disruption and leaky gut phenotype observed in larval intestines. Reduced expression of ion transporters (Oatp58Dc, Nhe3) further supports compromised epithelial polarity and barrier maintenance. Together, these findings highlight how AMP-specific aging programs compromise both cell-intrinsic homeostasis and overall tissue architecture.

We observed pronounced architectural shifts in AMP clusters under aging conditions, characterized by significant changes in both cluster frequency and spatial organization. While *Pirk* and *ND42* (mitochondrial complex I) knockdown induced clear morphological alterations, Toll activation failed to elicit similar changes, highlighting the genotype-specific effects of aging on progenitor architecture. Quantitative morphometric analysis including assessments of cluster shape, aspect ratio, solidity, and eccentricity confirmed a broad disruption of organizational integrity. These defects suggest that accelerated aging impairs the collective behaviour and spatial coordination of AMPs, rather than simply altering progenitor abundance. Mechanistically, these irregularities likely stem from the suppression of cell-cell adhesion, junctional stability, and cytoskeletal programs, as evidenced by our transcriptomic data. We further hypothesize that aberrant cluster morphology may result from the defective formation of the peripheral cells that normally encase AMPs. Because peripheral cells act as a transient niche regulating AMP proliferation and require Notch signaling for specification [28], the diminished Notch activity observed in our aging models may impair peripheral cell development, leading to the poorly defined boundaries and irregular morphologies reported here. To our knowledge, age-associated remodeling of AMP cluster architecture has not been previously documented. However, a recent finding has shown disrupted progenitor clustering in the aging adult intestine and described it as an early life biomarker of aging [135]. In accordance with this finding, our data show that genetic perturbation of cellular aging programs is sufficient to alter the spatial organization of progenitors in early developmental stages, implying that cluster architectural defects can be developmentally programmed and later contribute to age-associated intestinal dysfunction.

The emergence of aging-like defects following larval AMP-specific genetic perturbations led us to investigate whether these early-life insults exert lasting effects on adult intestinal homeostasis. We found that accelerated aging in progenitors during development is sufficient to “imprint” long-term lineage defects that persist beyond metamorphosis [31]. Specifically, constitutive activation of the Toll and Imd pathways during larval stages resulted in a sustained EE cell bias in the adult midgut. Conversely, anti-aging interventions such as overexpression of *Foxo* or *Atg8a* preserved wild-type lineage balance and further increased EC abundance, suggesting enhanced epithelial maintenance and differentiation potential. These observations align with established models where age-related stress drives aberrant stem cell behavior and a shift toward secretory fates [27, 31, 136, 160, 161]. Collectively, our findings demonstrate that developmental perturbations in the AMP niche are not transient; rather, they establish a “memory” of aging that progressively compromises adult homeostasis. This positions early progenitor health as a critical determinant of lifelong intestinal integrity and suggests that targeting specific developmental windows may offer potent preventive strategies against age-related decline.

## Materials and methods

### *Drosophila* Genetics and Husbandry

All *Drosophila* stocks and genetic crosses were maintained on a standard cornmeal medium in a 25°C incubator with a 12-h dark and light cycle. *Canton-S* was used as a wild-type control. A tissue-specific *Gal4* driver was employed to activate UAS-linked transgenes. Fly stocks used in this study included *esg-Gal4*,*UAS-GFP/CyO* (Chr. II, gifted by Edan Foley, University of Alberta, Canada), UAS-*pirk* RNAi (Chr. II, RRID: BL_67011), UAS-*toll10b* (Chr. X, RRID: BDSC_58987), UAS-*nd42* RNAi (Chr. III, RRID: BDSC_28894), UAS-*atg8*-GFP (Chr. III, RRID: BDSC_51656), and UAS-*foxo*-GFP (Chr. III, RRID: BDSC_43633).

### Antibodies used

Primary antibodies used were mouse anti-Prospero (1:25, DSHB Cat# Prospero (MR1A), RRID: AB_528440), mouse anti-H2AV (1:20, UNC93-5.2.1, DSHB; RRID: AB_2618077), mouse anti-Cora (1:25, C566.9, DSHB; AB_1161642), mouse anti-Cora (1:25, C615.16, DSHB; AB_1161644), rabbit anti-P62/SQSTM1 (1:250, Proteintech, RRID: AB_10694431), mouse anti-Armadillo (1:20, N27A1, DSHB; AB_528089) and rabbit anti-H3P (1:200, Sigma-Aldrich Cat# 06-570, RRID: AB_310177). CellROX™ Deep Red reagent (Invitrogen, C10422) was used for ROS detection. Normal goat serum (NGS; HiMedia, RM10701) was used as the blocking agent. Alexa Fluor 568–conjugated secondary antibodies included goat anti-mouse (1:400, Invitrogen, RRID: AB_144696) and goat anti-rabbit (1:400, Invitrogen, RRID: AB_10563566).

### Methods

Materials and methods section is provided in detail in the supplementary information.

## Acknowledgements

We would like to thank the Digital Imaging Facility (DIF) and the Common Instrumentation Facilities (CIF) at ACTREC for all the support. We thank the Bloomington Drosophila Stock Center, Developmental Studies Hybridoma Bank and the fly community for fly stocks and antibodies. Authors acknowledge HPC support by DBT grant BT/PR40181/BTIS/137/15/2021. We are thankful to the Stem Cell and Tissue Homeostasis lab for useful input and discussions.

## Author contributions

Conceptualization: R.J.K.; Data curation: R.J.K., S.M., A.A.M., S.J.P.; Formal analysis: S.M., A.A.M., S.J.P., I.S., M.S., P.C., M.M.I.; Funding acquisition: R.J.K.; Investigation: S.M., A.A.M., S.J.P., I.S., M.S.; Methodology: S.M., A.A.M., S.J.P., I.S., M.S.; Project administration: R.J.K.; Resources: R.J.K.; Supervision: R.J.K.; Validation: S.M., A.A.M., S.J.P., I.S., M.S.; Visualization: R.J.K.; Writing – original draft: S.M., A.A.M., R.J.K.; Writing – review & editing: S.M., A.A.M., R.J.K.

## Funding

This study was funded by the Department of Biotechnology, Ministry of Science and Technology, India for the Har Gobind Khorana – Innovative Young Biotechnologist Award (no. BT/13/IYBA/2020/14) to R.J.K., Ramalingaswami Re-entry Fellowship from the Department of Biotechnology, Ministry of Science and Technology, India (BT/RLF/Re-entry/19/2020) to R.J.K., ACTREC Postdoc fellowship to S.M. and ACTREC-SRF to A.A.M. This work was also funded by a Basic and Translational Research in Cancer grant (no.1/3(7)/2020/TMC/R&D-II/8823 Dt.30.07.2021), Capacity Building and Development of Novel and Cutting-edge Research Activities (no.1/3(4)/2021/TMC/R&D-II/15063 Dt.15.12.2021) from the Department of Atomic Energy, Government of India. M.S. is funded by CEFIPRA-CSRP (project no. 6704-5).

## Conflict of interest

The authors declare that there are no competing or financial interests

## Data availability statement

All the required data is within the article and supplementary information provided.

## Notes

### Competing Interest Statement

The authors have declared no competing interest.

