## Supplementary Information for "Developmental regulation of progenitor aging shapes long-term intestinal homeostasis in *Drosophila*"

#### Materials and Methods

##### Gut Dissection and Immunohistochemistry

Midguts from third instar larvae or adult flies were dissected in 1X Phosphate-buffered Saline (PBS) and fixed in a 1:1 mixture of 4% paraformaldehyde and n-heptane for 30 min at 28 °C. Following fixation, the lower aqueous phase was replaced with absolute methanol, and samples were gently inverted for 15-20 seconds. Samples were then washed three times in 1X PBS and permeabilized in 0.1% PBST (0.1% Triton X-100 in PBS) for 15 min.

Samples were blocked in 20% NGS for 20 min at room temperature and incubated with the respective primary antibody overnight at 4°C. After washing, samples were incubated with the appropriate secondary antibody for 2 hours at room temperature, followed by 1X PBS washes. Nuclei were stained with DAPI (2µg/ml) for 45 min, washed, and mounted in VECTASHIELD mounting medium (Vector Laboratories; RRID: AB_2336790).

##### Chemical treatment

Chemical treatment was performed using Paraquat or Rapamycin to modulate aging systemically. Early- to mid-third instar larvae from *esg-Gal4,UAS-GFP* x *wt* progeny were collected and placed in empty vials containing distilled water to prevent desiccation and starved for 2 h at room temperature. The larvae were then introduced in the test food containing Paraquat Dichloride Hydrate dissolved in water (20mM, Sigma Aldrich, 36541) along with its solvent control or Rapamycin dissolved in absolute ethanol (10µg/ml, Sigma Aldrich, R0395) along with its solvent control for 12 h. The treated mid- to late- 3$^{rd}$ instar larvae were dissected and used for further immunofluorescence-based experiments.

##### ROS detection

Reactive oxygen species were detected using the CellROX™ Deep Red reagent (Thermofisher scientific, C10422) following the manufacturer's protocol. Briefly, dissected, and fixed guts were washed in 1X PBS and incubated with 5µM CellROX reagent at 37 °C for 30 min before imaging by confocal microscopy.

##### SMURF assay

Intestinal barrier integrity was assessed using the SMURF assay. Second instar or early third instar larvae were starved for 4 h and transferred to food containing 2.5% (v/v) Brilliant Blue dye (Sigma Aldrich, 80717) for 18 h. Larvae were subsequently washed and scored as SMURF (blue dye diffusing outside the gut) or non-SMURF (blue dye contained within the gut).

For chemically treated larvae, Brilliant Blue dye was combined with 20mM paraquat or 10µg/ml rapamycin, and fed for 18 h post 4 h of starvation. The larvae were then scored phenotypically as mentioned above.

**Image processing, acquisition, and analysis**

All images were acquired using Zeiss LSM 780 or Zeiss LSM 980 microscopes using a 40x oil immersion objective (NA 1.4) at 0.6x zoom, unless otherwise mentioned. Images were quantified using ImageJ/Fiji (v1.54d, RRID: SCR_003070).

The raw image files were processed in ImageJ/Fiji software and converted to TIFF format (RGB mode), with individual or merged channels generated as required. TIFF files were subsequently imported into Adobe Photoshop (Creative Cloud) and assembled into figure panels on a RGB canvas. Final figures were saved in TIFF format using LZW compression at a resolution of 600 dpi or more.

**Image analysis:**

**Estimation of ROS and autophagy levels**

For reactive oxygen species (ROS) estimation in the midguts, the images for all the genotypes were acquired using the same settings/parameters for imaging. Quantification was performed using ImageJ/Fiji software. Mean fluorescence intensity (expressed in arbitrary units) was calculated for the midgut region after applying a uniform fluorescence threshold, which was defined based on the wild-type control and kept constant across all genotypes.

Autophagic flux in *esg* clusters was assessed by quantifying the total number of p62-positive puncta (p) and the total number of nuclei (n) within each cluster. The p62 puncta-to-cell ratio (p/n) was calculated assuming uninucleate *esg* clusters, where higher ratios indicate reduced autophagy flux (p62 accumulation) and lower ratios reflect enhanced autophagic degradation. The ratio of puncta per cell (p/n) was calculated, assuming all cells within the *esg* cluster to be uninucleate. This ratio represents the average number of autophagic puncta per cell and was used for comparison across different experimental genotypes.

**Counts of H3P and Prospero-positive cells in the midgut**

The total number of phospho-histone H3 (H3P)–positive and enteroendocrine (Prospero-positive) cells present throughout the midgut were manually counted from confocal images. Absolute numbers of H3P and Prospero-positive cells per midgut were plotted using GraphPad Prism version 9.0.

**Quantification of enterocytes in the adult midgut**

The total number of enterocytes present in the adult midgut were quantified by counting DAPI nuclei in the whole midgut region.

**Quantification of H2Av-positive cells in the midgut:**

DNA damage in esg-GFP–positive cells was assessed by immunostaining for H2Av. The number of H2Av-positive cells within the esg-expressing population was manually counted from confocal images and normalized to the total number of esg-positive cells per midgut. The percentage of DNA damage was calculated as the proportion of H2Av-positive esg cells relative to the total esg cell population and multiplied by 100.

**Analysis of Coracle levels:**

To assess the effect of aging on the septate junction protein Coracle, fluorescence intensity was measured using the line intensity measurement tool in ImageJ software. The average intensity per gut was calculated and plotted.

**Quantification of GFP-Positive Cluster Number and Morphometric Analysis:**

**Image Acquisition and Cluster Counting**

Two representative images per larva (n ≥ 12 larvae per genotype) were analyzed to quantify the total absolute number of GFP-positive clusters for each genotype.

**Image Processing and Morphometric Analysis**

**Image Pre-processing and Segmentation:**

To facilitate high-throughput analysis, three-dimensional Z-stacks were first reduced to two-dimensional images. This was achieved by applying a Maximum Intensity Z-projection in ImageJ/Fiji. Given the relatively planar morphology of the cell clusters, this 2D projection served as a reliable approximation, significantly reducing the computational demand for subsequent segmentation and analysis.

Initial segmentation of the cell clusters was performed using the interactive machine learning software Ilastik (v1.4.0). A pixel classification model was trained on a representative subset of the projected images to distinguish between cluster and background pixels. The trained classifier was then applied in a batch process to automatically segment the entire experimental dataset, generating binary masks for each projected image. Each cluster was assigned a unique label ID for further analysis.

**Manual Curation and Post-processing:**

The raw images and their corresponding segmentation masks were imported into the Napari multi-dimensional image viewer for Python for manual review and correction. This curation step was critical to ensure the accuracy of the final quantifications. Manual corrections included: i) Refining the boundaries of segmented clusters to more accurately match the source image. ii) Separating distinct but touching clusters that were erroneously merged during automated segmentation. iii) Excluding clusters that were partially captured at the image frame boundary, incompletely segmented due to low signal intensity, or otherwise artifactual.

**Cluster Quantification:**

Before feature extraction, the intensity values of the raw grayscale images were normalized to a range of [0, 1]. To avoid bias from incomplete objects, clusters touching the borders of the image were programmatically removed using the `pyclesperanto_prototype.exclude_labels_on_edges()` function. A comprehensive set of morphometric features for each cluster was quantified from the curated masks using the `skimage.measure.regionprops_table()` function within the scikit-image library in Python.

**Description of Quantified Morphometric Properties**

The following morphometric and intensity parameters were extracted for each segmented cluster using the `skimage.measure.regionprops_table()` function: i) Label: Unique integer identifier assigned to each segmented cluster. ii) Area: Total number of pixels within the cluster. iii) Centroid: (Row, column) coordinates of the cluster's geometric center. iv) Feret diameter max: The longest distance between any two points along the boundary of the cluster. v) Solidity: Ratio of the cluster's area to the area of its convex hull. This is a measure of the cluster's convexity, with a value of 1 indicating a perfectly convex shape. vi) Eccentricity: Measure of deviation from circularity, where 0 corresponds to a circle and 1 to a line segment. vii) Axis major length: Length of the major axis of the ellipse that has the same normalized second central moments as the cluster. viii) Axis minor length: Length of the minor axis of the ellipse that has the same normalized second central moments as the cluster. ix) Area convex: Number of pixels in the convex hull of the cluster. x) Perimeter: Length of the cluster boundary. xi) Intensity std: The standard deviation of the intensity values of the pixels within the cluster. xii) Intensity mean: The average intensity value of the pixels within the cluster.

**Transcriptomics Analysis:**

**RNA Isolation and Quality Assessment**

For each biological replicate, ~1µg of RNA sample was isolated from dissected larval midguts and outsourced to Medgenome for library preparation and sequencing. Briefly, larval midguts were dissected in cold 1X PBS, and lysed in 500µl of Trizol Reagent (Ambion–Life Technologies, 11596018). Total RNA was extracted using the phenol-chloroform method, and RNA yield and quality were quantified using Nanodrop spectrophotometer.

**Library Preparation and Sequencing**

RNA sequencing library preparation and sequencing were carried out by MedGenome Labs Ltd. Libraries were generated using a strand-specific RNA-seq protocol based on the NEBNext Ultra II Directional RNA Library Prep Kit with minor modifications. Polyadenylated RNA was enriched from total RNA using oligo(dT)-coupled magnetic beads. The purified mRNA was subsequently fragmented under elevated temperature in the presence of divalent cations.

Fragmented RNA was reverse transcribed using random hexamer primers to synthesize first-strand cDNA, followed by second-strand synthesis incorporating uracil in place of thymine to enable strand specificity. Strand directionality was preserved by selective digestion of the uracil-containing second strand using the USER (Uracil-specific Excision Reagent) enzyme treatment. The resulting single-stranded cDNA fragments were indexed and amplified by limited-cycle PCR, followed by AMPure XP bead purification to obtain sequencing-ready libraries.

Final libraries were sequenced on an Illumina HiSeq X or NovaSeq platform to generate 2 × 150 bp paired-end reads per sample. Sequencing quality was ensured, with up to 75% of bases achieving a Phred quality score of Q30 or higher. Raw sequencing output was demultiplexed to generate FASTQ files, which were provided via secure FTP for downstream bioinformatic analysis.

**Data Analysis:**

**Gene Expression Analysis**

The quality check (QC) for raw FASTQ files was performed using FASTQC (v0.12.1). Low-quality reads and adapter sequences were trimmed using FASTP (v0.23.4) [1] and further followed by a QC using FASTQC. The good-quality FASTQ reads were further aligned to the *Drosophila* BDGP6 transcriptome using Salmon in quasi-mapping mode, enabling transcript abundance quantification with GC bias and sequence-level bias corrections [2]. The transcript-level abundances were converted to gene-level counts using the tximport package available in R [3]. Differential gene expression analysis was performed using DESeq2, with adjusted p-value (padj)& lt;0.05 [4]. The differentially expressed genes were visualized using a volcano plot and heat maps.

**Functional Enrichment Analysis**

The Gene Set Enrichment Analysis was performed using Webgestault (WEB-based GEne SeT AnaLysis Toolkit) [5] with log fold changes obtained from DESeq2 analysis, utilizing databases like KEGG (Kyoto encyclopedia of genes and genomes)[6], Reactome[7], Panther [8], and Wikipathways [9]. For each database, pathways with normalized enrichment scores $< -1$ or $> 1$ were filtered. Barplot representing the hallmark aging pathways enriched in the analysis for each database. Customized Python scripts were created to parse Webgestault outputs to obtain pathway-wise enriched genes. Heatmaps were generated to visualize the enriched gene expression patterns using DESeq2 normalized counts.

**Statistical Analysis**

Statistical analysis was performed using the GraphPad Prism Version 8.0.1 software (RRID: SCR_002798). For analysis of statistical significance, each experimental sample was tested with its respective control in each experimental setup for all the data in each of the figures to estimate the *P*-value. Statistical significance was determined by using Welch's unpaired Student's *t*-test. Data are presented as mean ± SD. Exact *p*-values are indicated in the graphs. Mutant genotypes were compared with wild-type controls, while knockdown or overexpression genotypes were compared with their respective parental Gal4 controls crossed to wild-type flies for all statistical analyses. Experiments involving chemical treatments were compared with their corresponding vehicle controls. No statistical methods were used to predetermine sample size, and experiments were not randomized.

**Supplementary figure legends:**

**Figure S1: Genetic modulation of cellular aging in AMPs regulates DNA damage accumulation.**

Representative images (A–F'') of γH2AX staining in AMPs upon *esg-Gal4* specific expression of *UAS-toll10b, UAS-pirkRNAi, UAS-nd42RNAi, UAS-foxoGFP,* or *UAS-atg8aGFP* as compared to wild-type control. Arrowheads indicate γH2AX-positive nuclei (in the Escargot-GFP positive islets). Quantitation of the percentage of γH2AX-positive esg$^+$ progenitor cells in the indicated genotypes relative to wild-type control (G). Boxed regions in panels (A-F) are shown as magnified images in the corresponding panels (A'-F', A''-F''). GFP expression (green) is driven by *esg-Gal4,UAS GFP* and nuclei stained by DAPI (blue). Scale bar: 20 µm (A-F), 5 µm (A'-F', A''-F''). A minimum of 10 larvae were considered and each data point represents γH2AX-positive esg cells with respect to total esg cells. Statistical significance was assessed using Student's t-test with Welch's correction. P-values are indicated in the respective graphs.

**Figure S2: Chemical intervention-based modulation of aging regulates septate junctions and gut barrier integrity**.

Schematic illustrating the experimental workflow for Paraquat (20mM) or Rapamycin (10µg/ml) treatment of *Drosophila* larvae (A). Representative images of Coracle (Red) staining in larval midguts following Paraquat (C–C') or Rapamycin (E-E') treatment relative to vehicle control (B-B', D-D'). Quantitation of mean fluorescence intensity of Coracle upon Paraquat (F) or Rapamycin (H) treatment. Representative Smurf assay images in larvae following Paraquat (G) and Rapamycin (I) treatment. GFP expression (green) is driven by *esg-Gal4,UAS GFP* and nuclei stained by DAPI (blue). Boxed regions in panels (B-E) are shown as magnified images in the corresponding panels (B'–E'). A minimum of 8 larvae were considered for Coracle quantitation. Each data point represents the mean fluorescence intensity calculated from 5 individual measurements per image. For the SMURF assay, a minimum of 8 larvae were considered. Scale bar: 20 µm (B-E), 5 µm (B'-E'). Statistical significance was assessed using Student's t-test with Welch's correction. P-values are as indicated in the graphs. Schematic (A): Created in BioRender. Khadilkar, R. (2026) https://BioRender.com/4lz1fnv.

**Fig. S3: Chemical intervention-based modulation of aging regulates DNA damage accumulation in the midgut**

Schematic illustrating the experimental workflow for Paraquat (20mM) or Rapamycin (10µg/ml) treatment of *Drosophila* larvae (A). Representative images of vehicle control (B-B''), Paraquat-treated (C-C'') or Rapamycin treated (D-D'') larval intestines stained for γH2AX. Arrowheads indicate γH2AX-positive nuclei. Quantitation of the percentage of γH2AX-positive *esg*$^+$ cells following Paraquat and Rapamycin treatments (F–G). GFP expression (green) is driven by *esg-Gal4,UAS GFP* and nuclei stained by DAPI (blue). Boxed regions in panels (B-E) are shown as magnified images in the corresponding panels (B'–E',

B''-E''). Scale bar: 20 µm (B-E), 5 µm (B'-E'). A minimum of 10 larvae were considered and each data point represents γH2AX-positive esg cells with respect to total esg cells. Statistical significance was assessed using Student's t-test with Welch's correction. P-values are as indicated in the graphs. Schematic (A): Created in BioRender. Khadilkar, R. (2026) https://BioRender.com/8l9jk6o.

**Figure S4: Differential gene expression and pathway enrichment analyses highlight aging-associated transcriptional changes in AMP- specific *pirk*-knockdown versus Foxo-overexpressing midguts**

Principal Component Analysis (PCA) plot for biological replicates of *Pirk* knockdown (test) and Foxo overexpressed (reference) genotype gut cells (A). Volcano plot representing significantly deregulated genes (B). Barplot representing pathway enrichment scores obtained from Gene Set Enrichment Analysis performed with KEGG, Panther, Reactome, and Wikipathways database (C). Heatmap representing differentially expressed genes involved in hallmark aging pathways (D). Heatmap representing overall differentially expressed genes (E).

**Figure S5: Differential gene expression and pathway enrichment analyses highlight aging-associated transcriptional changes in AMP-specific ND42 knockdown versus Foxo-overexpressing midguts**

Principal Component Analysis (PCA) plot for biological replicates of ND42 Knockdown (test) and Foxo overexpressed (reference) genotype gut cells (A). Volcano plot representing significantly deregulated genes (B). Barplot representing pathway enrichment scores obtained from Gene Set Enrichment Analysis performed with KEGG, Panther, Reactome, and Wikipathways database (C). Heatmap representing differentially expressed genes involved in hallmark aging pathways (D). Heatmap representing overall differentially expressed genes (E).

**Figure S6: Genetic modulation of cellular aging in larval AMPs has a developmental impact on enterocyte numbers in the adult midgut**

Schematic illustrating the impact of AMP-specific genetic modulation on adult gut homeostasis (A). Representative images (B–F) showing enterocyte populations in adult midguts, starting from wild-type (B), *UAS-toll10b* (C), *UAS-pirkRNAi* (D), *UAS-foxoGFP* (E) or *UAS-atg8aGFP* (F), aging genotypes (C–D), and anti-aging genotypes (E–F). Quantitation of enterocyte number per adult midgut across genotypes (G). Each data point represents the number of enterocytes per midgut. Nuclei stained by DAPI (magenta). A minimum of 10 adults were used for the experiment. Scale bar: 20 µm (B-F). Statistical significance was assessed using Student's t-test with Welch's correction. P-values are as indicated in the graphs. Schematic (A): Created in BioRender. Khadilkar, R. (2026) https://BioRender.com/ebtc42r.

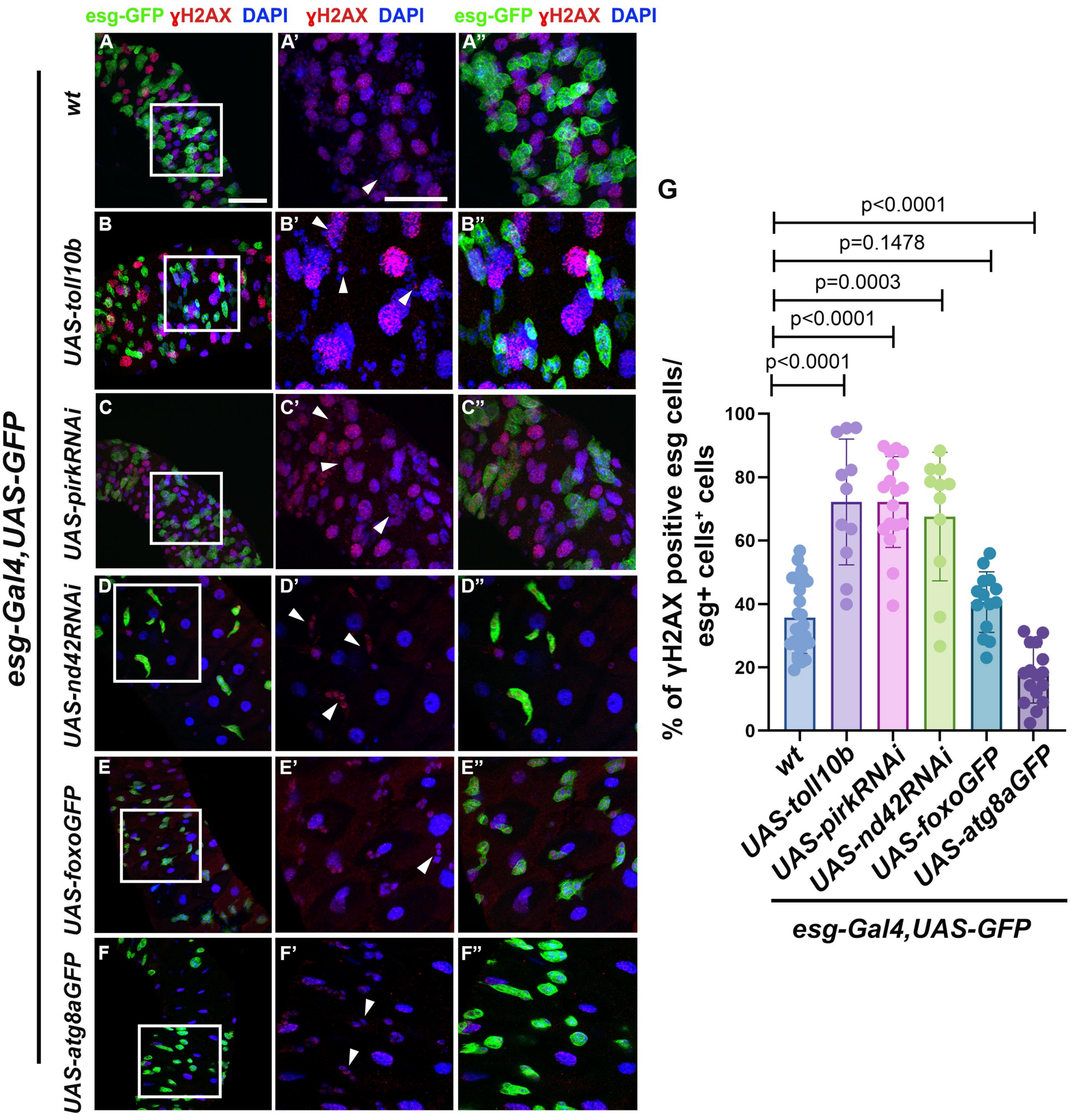
esg-GFP ɣH2AX DAPI
ɣH2AX DAPI
esg-GFP ɣH2AX DAPI
A
A'
A''
B
B'
B''
C
C'
C''
D
D'
D''
E
E'
E''
F
F'
F''
wt
UAS-toll10b
UAS-pirkRNAi
UAS-nd42RNAi
UAS-foxoGFP
UAS-atg8aGFP
esg-Gal4,UAS-GFP
G
p<0.0001
p=0.1478
p=0.0003
p<0.0001
p<0.0001
% of γH2AX positive esg cells/ esg+ cells⁺ cells
100
80
60
40
20
0
wt
UAS-toll10b
UAS-pirkRNAi
UAS-nd42RNAi
UAS-foxoGFP
UAS-atg8aGFP
esg-Gal4,UAS-GFP

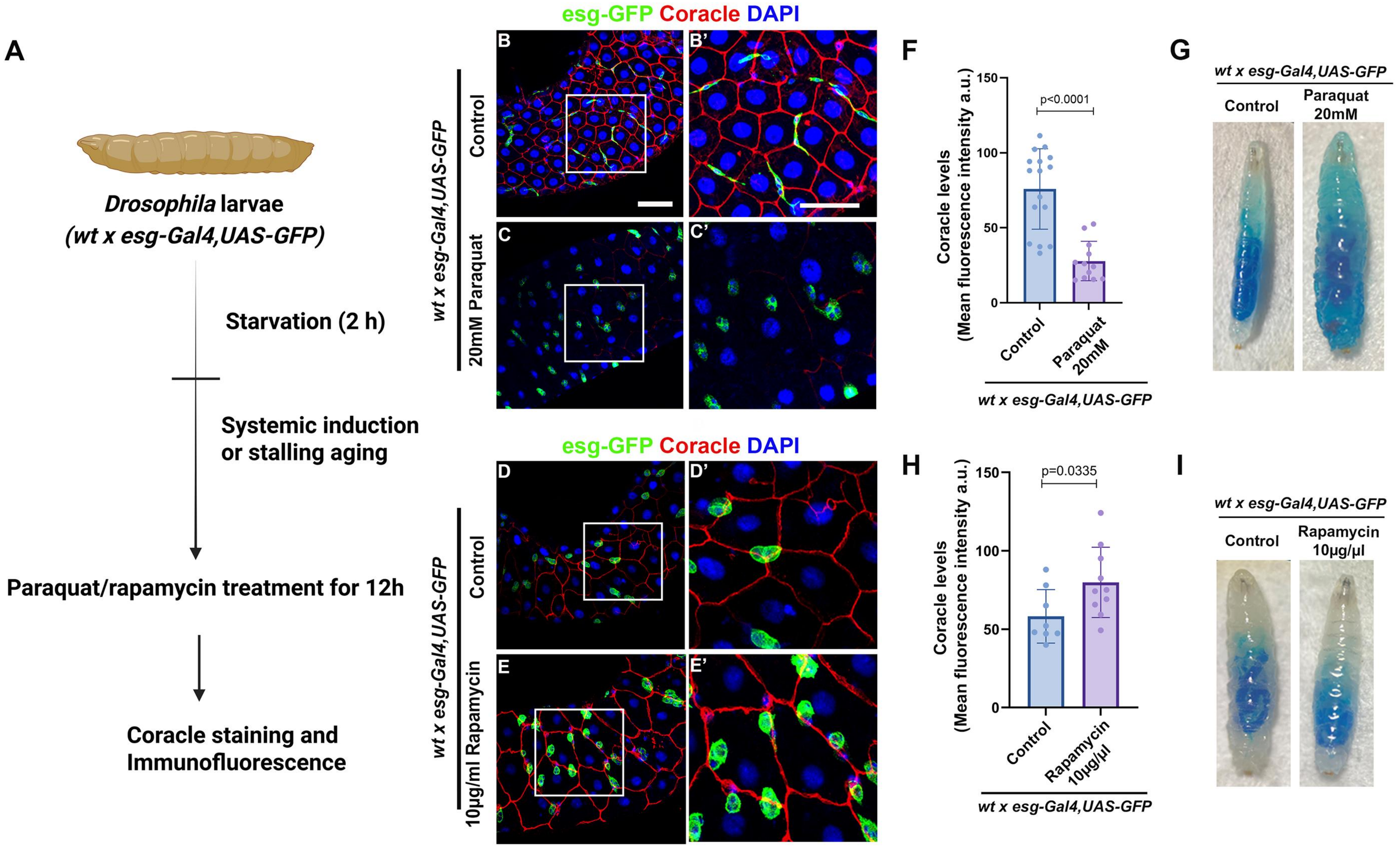
A
Drosophila larvae
(wt x esg-Gal4,UAS-GFP)
Starvation (2 h)
Systemic induction
or stalling aging
Paraquat/rapamycin treatment for 12h
Coracle staining and
Immunofluorescence
esg-GFP Coracle DAPI
B
B'
C
C'
wt x esg-Gal4,UAS-GFP
Control
20mM Paraquat
D
D'
E
E'
Control
10μg/ml Rapamycin
F
Coracle levels
(Mean fluorescence intensity a.u.)
p<0.0001
Control
Paraquat
20mM
wt x esg-Gal4,UAS-GFP
G
wt x esg-Gal4,UAS-GFP
Control
Paraquat
20mM
H
p=0.0335
Control
Rapamycin
10μg/μl
I
Control
Rapamycin
10μg/μl

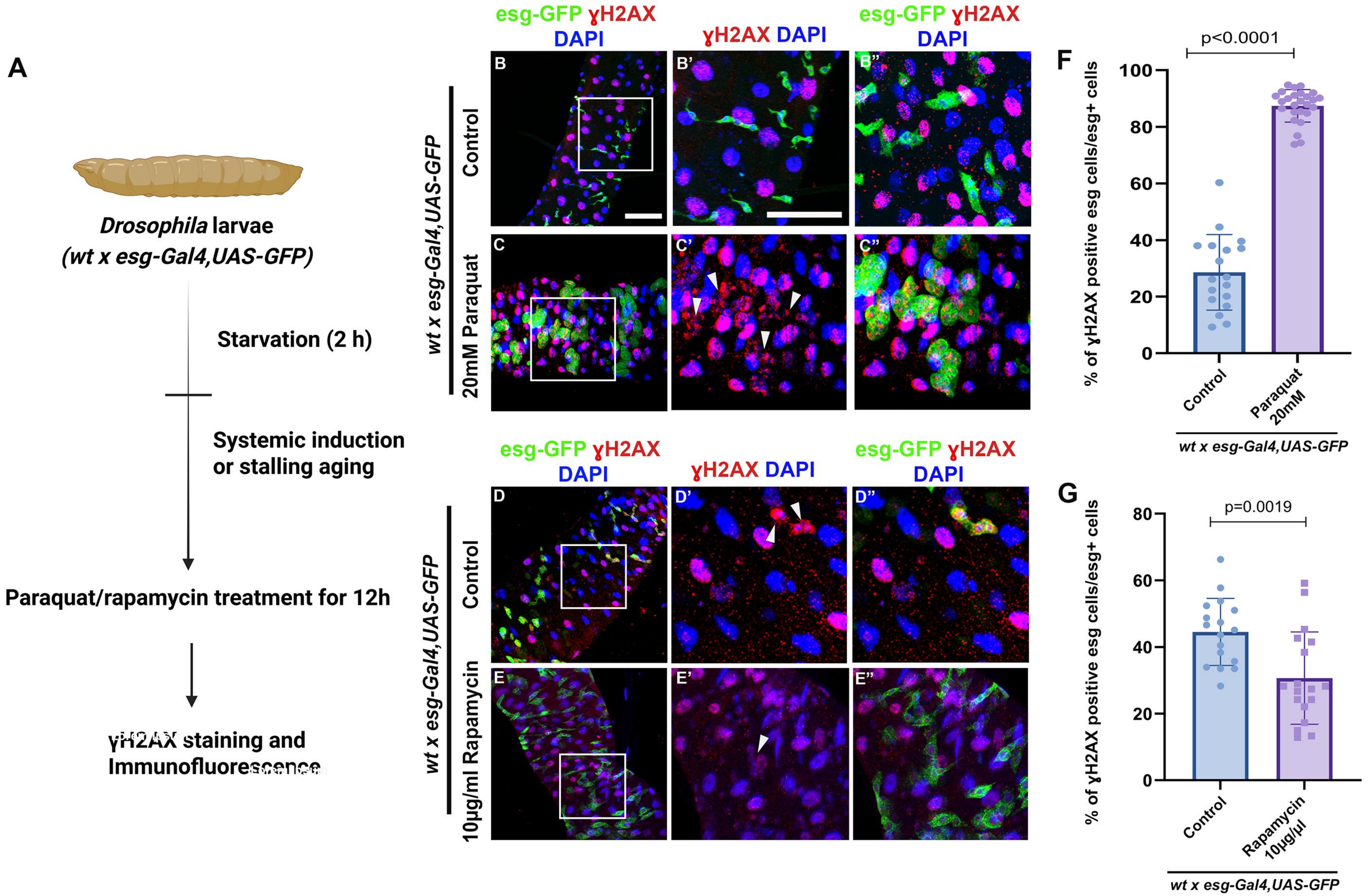
A
Drosophila larvae
(wt x esg-Gal4,UAS-GFP)
Starvation (2 h)
Systemic induction
or stalling aging
Paraquat/rapamycin treatment for 12h
γH2AX staining and
Immunofluorescence
esg-GFP γH2AX DAPI
γH2AX DAPI
esg-GFP γH2AX DAPI
wt x esg-Gal4,UAS-GFP
Control
20mM Paraquat
B
B'
B''
C
C'
C''
D
D'
D''
E
E'
E''
10µg/ml Rapamycin
F
p<0.0001
% of γH2AX positive esg cells/esg+ cells
100
80
60
40
20
0
Control
Paraquat 20mM
wt x esg-Gal4,UAS-GFP
G
p=0.0019
80
60
40
20
0
Control
Rapamycin 10µg/µl
wt x esg-Gal4,UAS-GFP

### Aging (Pirk) v/s antiaging (Foxo)

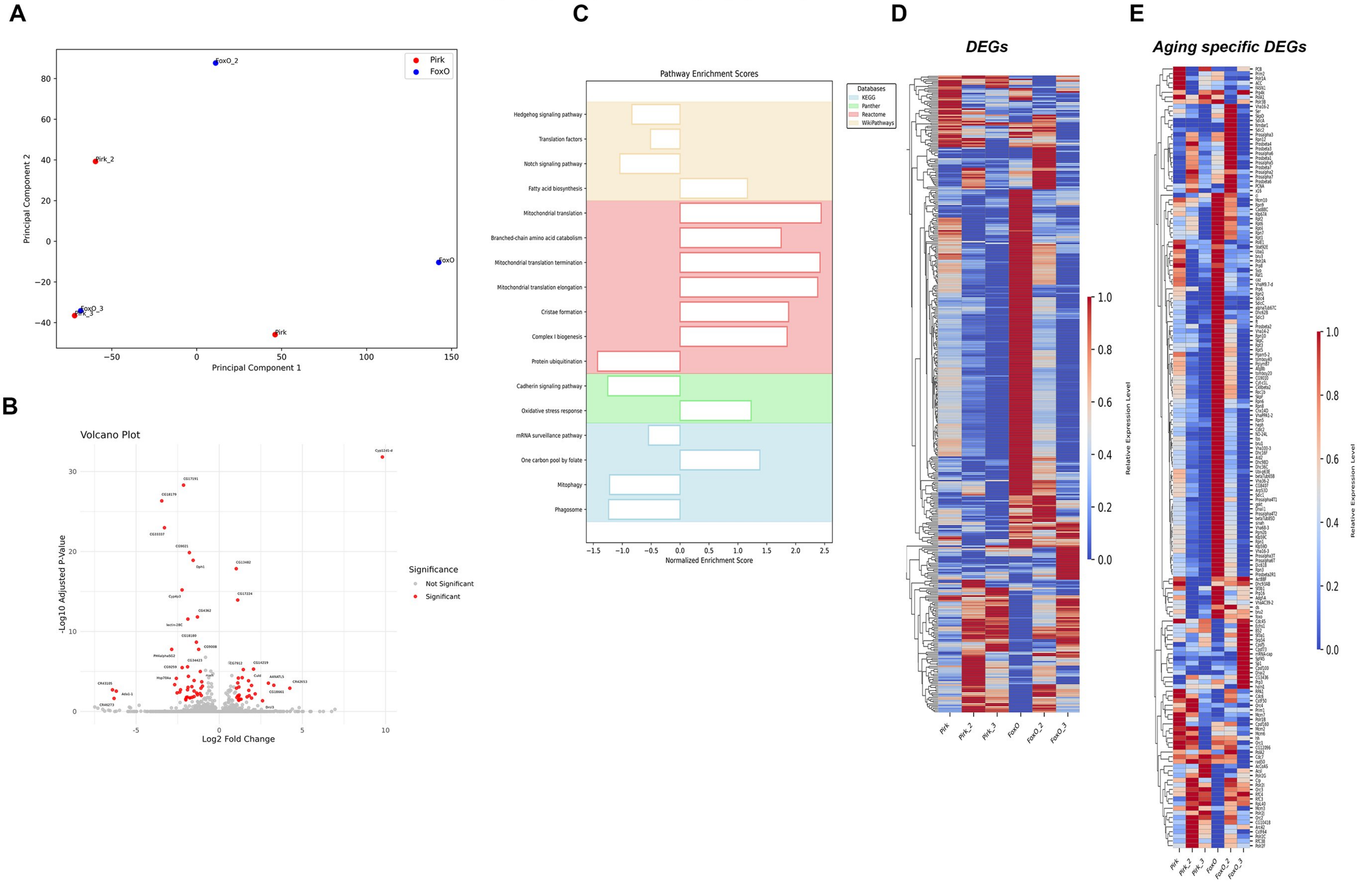

### Aging (ND42) v/s antiaging (Foxo)

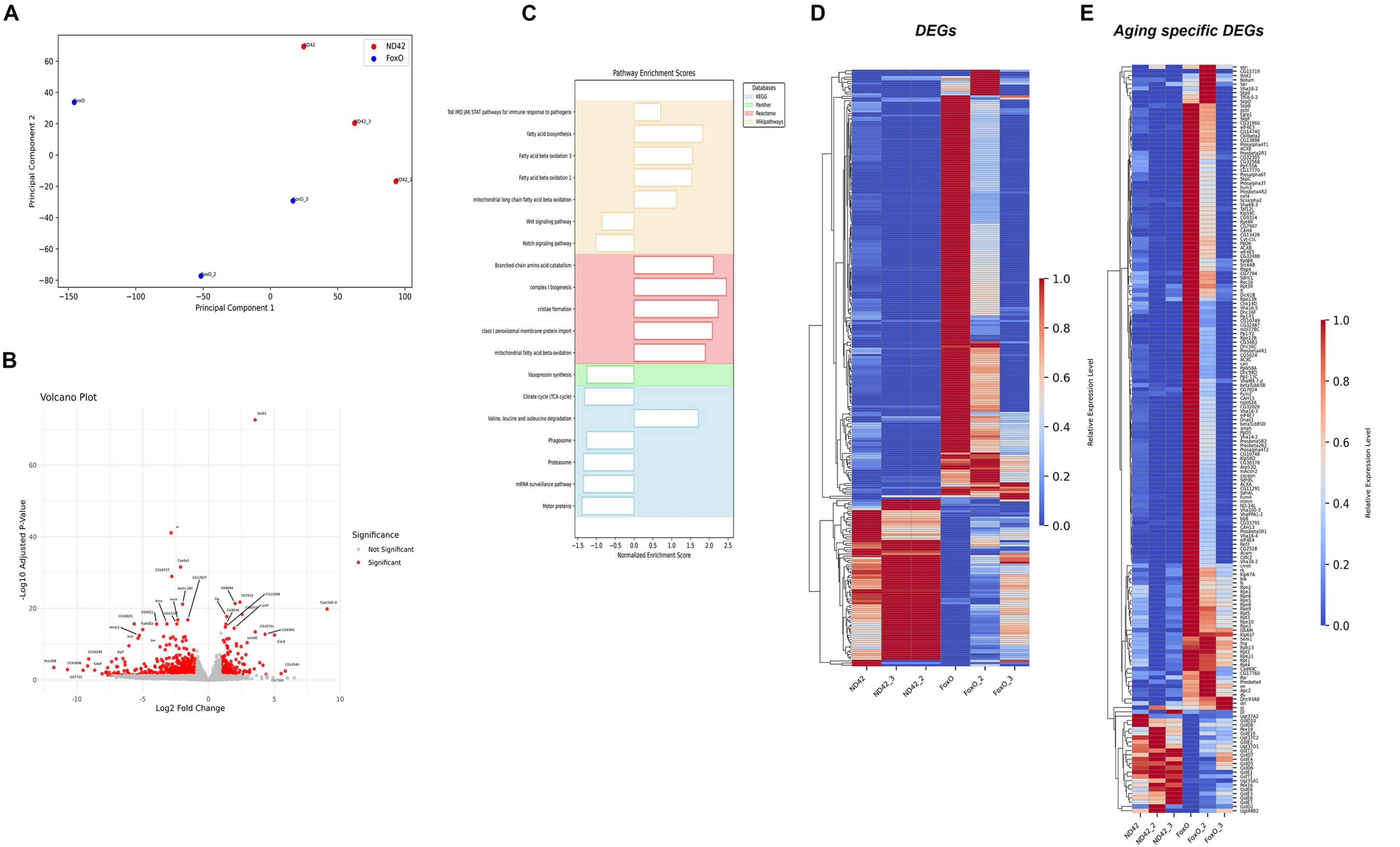

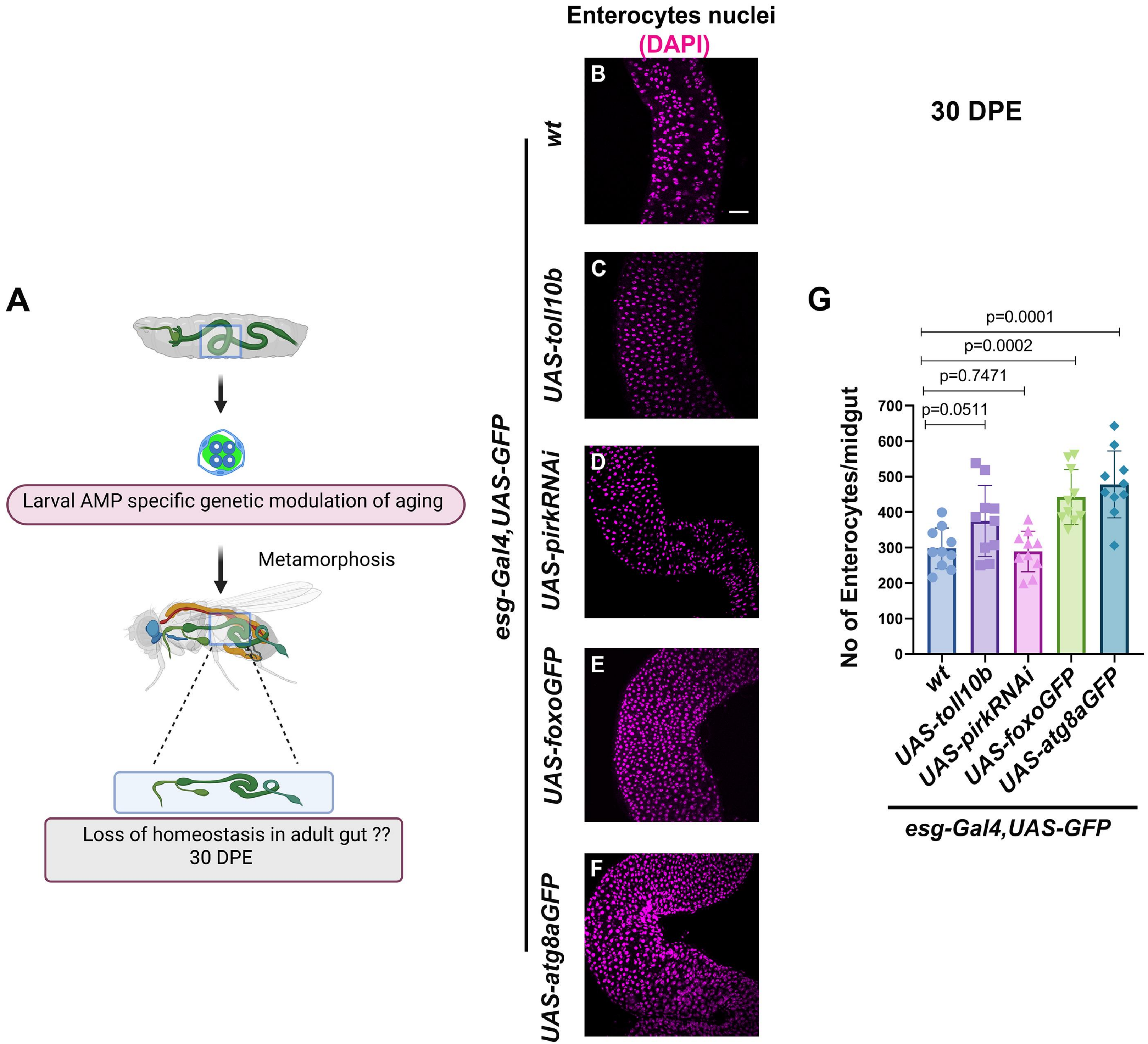
A
Larval AMP specific genetic modulation of aging
Metamorphosis
Loss of homeostasis in adult gut ??
30 DPE
Enterocytes nuclei
(DAPI)
esg-Gal4,UAS-GFP
wt
UAS-toll10b
UAS-pirkRNAi
UAS-foxoGFP
UAS-atg8aGFP
B
C
D
E
F
30 DPE
G
p=0.0001
p=0.0002
p=0.7471
p=0.0511
No of Enterocytes/midgut
0
100
200
300
400
500
600
700
wt
UAS-toll10b
UAS-pirkRNAi
UAS-foxoGFP
UAS-atg8aGFP
esg-Gal4,UAS-GFP
